# A vast repertoire of secondary metabolites influences community dynamics and biogeochemical processes in cold seeps

**DOI:** 10.1101/2023.08.12.552926

**Authors:** Xiyang Dong, Tianxueyu Zhang, Weichao Wu, Yongyi Peng, Xinyue Liu, Yingchun Han, Xiangwei Chen, Zhizeng Gao, Jinmei Xia, Zongze Shao, Chris Greening

## Abstract

In deep sea cold seeps, diverse microbial communities thrive on the geological seepage of hydrocarbons and inorganic compounds. These chemosynthetically-driven communities are unique in composition, ecology, and biogeochemical activities compared to photosynthetically-driven ecosystems. However, their biosynthetic capabilities remain largely unexplored. Here, we analyzed 81 metagenomes, 33 metatranscriptomes, and seven metabolomes derived from nine globally distributed areas of cold seeps to investigate the secondary metabolites produced by cold seep microbiomes. Cold seep microbiomes encode diverse, abundant, and novel biosynthetic gene clusters (BGCs). Most BGCs are affiliated with understudied bacteria and archaea, including key mediators of methane and sulfur cycling, and multiple candidate phyla. The BGCs encode diverse antimicrobial compounds (e.g. NRPS, PKSs, RiPPs) that potentially shape community dynamics, as well as compounds predicted to influence biogeochemical cycling, such as phosphonates, iron-acquiring siderophores, nitrogenase-protecting glycolipids, and methyl-CoM reductase-modifying proteins. BGCs from key players in cold seeps are widely distributed and highly expressed, with their abundance and expression levels varying with different sediment depths. Numerous unique natural products were detected through untargeted sediment metabolomics, demonstrating a vast, unexplored chemical space and validating *in situ* expression of the BGCs in cold seep sediments. Overall, these results demonstrate cold seep sediments potentially serve as a reservoir of hidden natural products and provide insights into microbial adaptation in chemosynthetically-driven ecosystems.

## Introduction

Deep sea cold seeps are unique and extraordinary ecosystems primarily located at continental margins. These seepages originate from the upward migration of hydrocarbon-rich fluids (mainly methane) through submarine microfractures^1, 2^. Chemosynthetic microbial consortia transform these geologically-derived hydrocarbons and inorganic compounds into biomass. Particularly dominant are anaerobic methane-oxidizing archaea (ANME) that live syntrophically with sulfate-reducing bacteria (SRB)^3^. Due to these activities, cold seeps are hotspots of biomass and diversity in the deep sea^4^, harbouring compositionally and functionally unique microbial communities^5, 6^. Nevertheless, extensive competition for resources and space in these sediments exerts strong eco-evolutionary pressures^7–9^. Thus, microbes employ a plethora of strategies to adapt to these unique environments.

Microorganisms encode biosynthetic gene clusters (BGCs) to synthesize natural products (NPs), also known as secondary or specialised metabolites (SMs). Microbes collectively produce an extraordinary array of NPs, which vary in their chemical structures, biosynthesis pathways, and physiological functions, many of which have pharmaceutical applications^10–12^. The major classes produced include ribosomally synthesized and post-translationally modified peptides (RiPPs), polyketide synthases (PKSs), non-ribosomal synthetic peptides (NRPS), PKS-NRPS hybrids, and terpenes. Many NPs have antimicrobial properties: microbes employ them as biological weapons^13–15^ to kill or inhibit competition, protect the host cells from predators and pathogens^16^, and gain a competitive advantage in nutrients and space^17^. However, NPs also enable microbes to adapt to their environments in other ways. For example, siderophores enhance iron transport and bioavailability in cells^18^, aryl polyenes (APEs) covalently attached in biofilms protect cells from damage caused by reactive oxygen species (ROS)^19^, and compatible solutes protect against osmotic stress^20^. In addition to facilitating environmental adaptation of microbes, NPs more broadly contribute to the evolutionary pressures and ecological dynamics shaping ecosystems^21–23^.

Comparative genomics data indicate that numerous unknown and hidden NPs remain to be discovered in uncultivated bacteria and archaea^24–26^. Several tools are available to analyze the secondary metabolic potential of these uncultured microorganisms, such as antiSMASH^27^, DeepBGC^28^, SanntiS^29^, and GECCO^30^; these identify BGCs by searching homologues of core biosynthetic genes or via utilization of deep learning and natural language processing (NLP) strategy^29, 31, 32^. Seminal studies in this area focused on soil ecosystems and emphasized that both well-known antimicrobial producers (i.e. Actinobacteriota) and understudied phyla (e.g. Acidobacteriota and Verrucomicrobiota) dominate NP biosynthesis^24, 31^. More recent studies have examined the production of bioactive NPs by the microbiota of plants and animals in recent years, such as in the human and animal digestive tracts^33, 34^, plant rhizospheres^35^ and marine sponges^36^, and highlighted their importance in regulating health and disease within their hosts. BGC repertoires have been explored in numerous other environments, spanning global seawaters^37, 38^, anoxic basins^39^, coastal marine sediments^40^, Antarctic deserts^41^, and glaciers^42^, as well as engineered biomes^43^. However, the secondary metabolite potential of the deep biosphere and chemosynthetically-driven ecosystems such as cold seeps has largely been overlooked. Most NP research has also focused on bacteria, with minimal ecosystem-level research on the potential of archaea to produce secondary metabolites^44^.

In this study, we addressed these knowledge gaps by performing an in-depth analysis of the BGCs encoded by microorganisms and their synthesized NPs in global cold seeps. To do so, we analyzed metagenomics, metatranscriptomics, and metabolomics datasets from nine cold seeps **(Figure 1 and Supplementary Table 1)**. By integrating these data, we identified a wide variety of microbial BGCs from 2,479 metagenome-assembled genomes (MAGs), including multiple uncultivated (candidate) bacterial and archaeal phyla. We also detected numerous novel NPs through mass spectrometry. These findings enhance understanding of the adaptive mechanisms and interspecies competition of microbial communities in the deep biosphere. Moreover, they provide a pathway to identify new antimicrobial compounds and other types of drugs.

**Figure 1.**
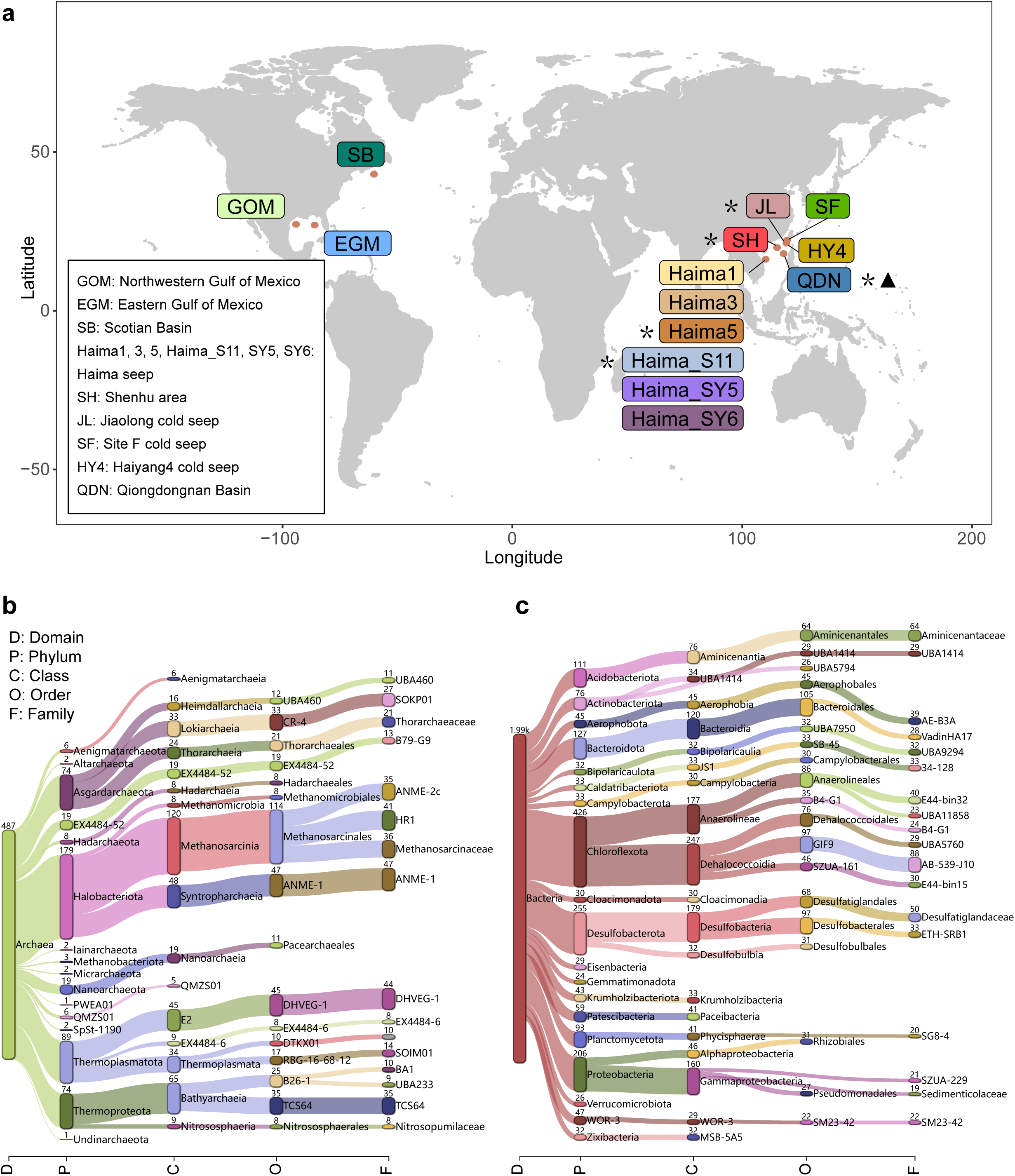
The map of nine globally distributed cold seep sites and reconstruction of MAGs. **(a)** Geographic distribution of all cold seep sites where metagenomic data were collected. Asterisks donate sites where metatranscriptomes were also collected, the triangle denotes the site were metabolomes were collected. Details are shown in **Supplementary Table 1**. **(b)-(c**) Sanket plots showing recovered MAG information of cold seep sediment microbiome at different taxonomic levels based on GTDB-Tk classification, including **(b)** archaea and **(c)** bacteria. The numbers indicate the number of MAGs recovered for the lineage. Detailed trees for recovered MAGs can be found in **Supplementary Figure 1**.

## Results and Discussion

### Cold seep microbiomes harbor diverse and unique BGCs

We investigated the biosynthetic potential of the cold seep microbiome at global scale by first analyzing 2,479 previous constructed metagenome-assembled genomes (MAGs; 1,992 bacterial, 487 archaeal) spanning 89 phyla^7, 45^ (**Figure 1, Supplementary Figure 1 and Supplementary Table 2)**. A total of 2,865 BGCs (longer than 10 kb; 2,627 bacterial and 238 archaeal) were predicted **(Figure 2a, b and Supplementary Table 3)**. These BGCs encoded for seven of the eight BiG-SCAPE classes^46^: ribosomally synthesized and post-translationally modified peptides (RiPPs, n = 580), terpenes (n = 480), non-ribosomal peptides (NRPS, n = 444), type I polyketide synthases (PKSI, n = 117), other polyketide synthases (Other PKSs, n = 353), polyketide-non-ribosomal peptide combinations (PKS-NRPS hybrids, n = 19), and Others (n = 872). In line with other studies^39, 41, 43, 47^, multiple incomplete BGCs were retrieved at the end of metagenomic contigs.

**Figure 2.**
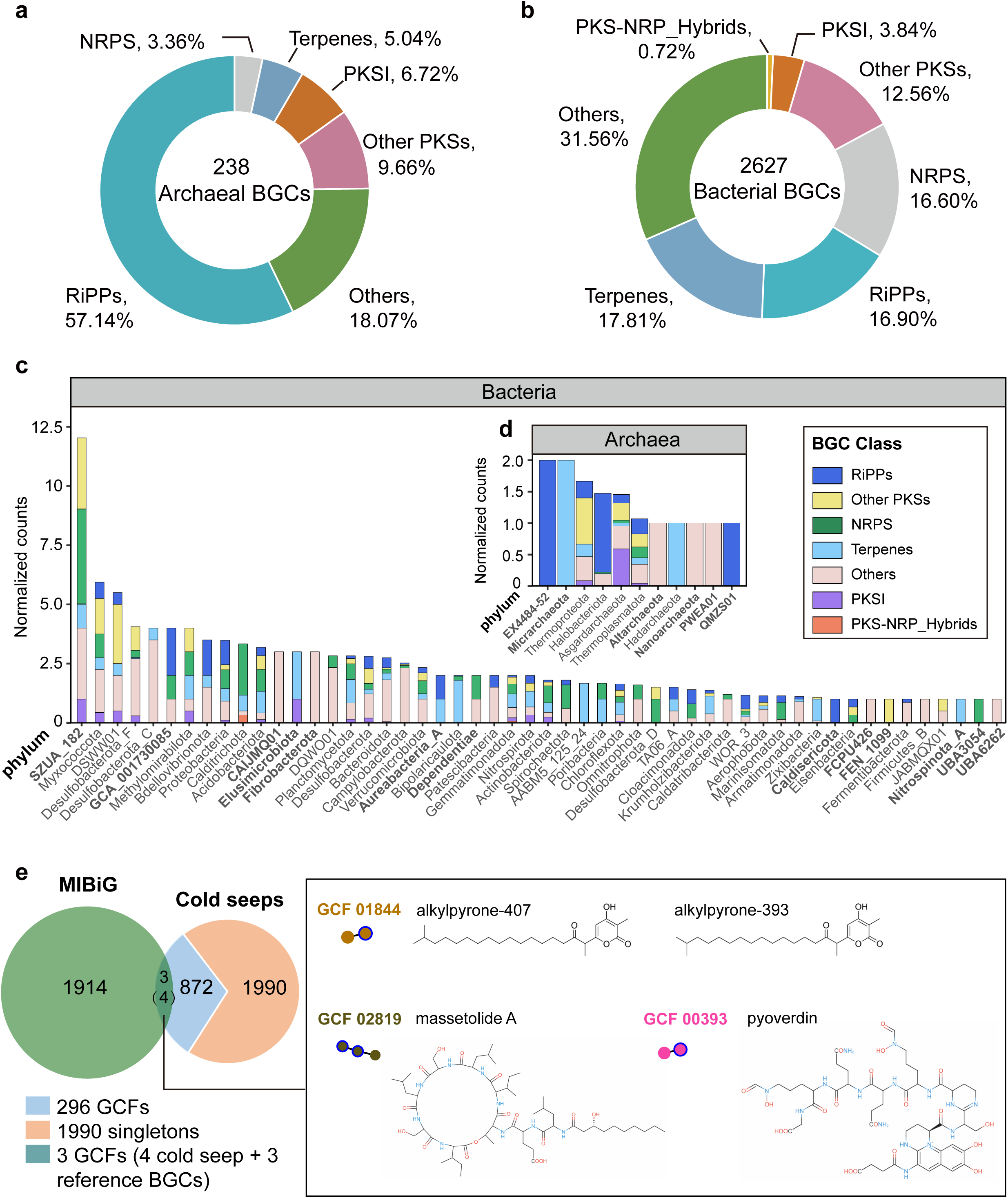
Biosynthetic gene clusters detected in cold seep archaeal and bacterial MAGs. Relative proportions of different BGC classes in (**a)** archaeal and (**b)** bacterial MAGs derived from 81 cold seep sediment samples based on BiG-SCAPE. **(c)** Normalized counts of bacterial biosynthetic gene clusters at the phylum level. **(d)** Normalized counts of archaeal biosynthetic gene cluster at the phylum level. Bold labels indicate phyla with only one representative MAG. Normalized BGC counts were derived by dividing the total count of each BGC type present in a phylum by the total number of MAGs from that phylum. **(e)** Venn diagram showing the overlap between the MIBiG database and cold seep BGCs. Three cold seep BGCs overlap with four reference BGCs, forming three Gene Cluster Families (GCFs). Detailed BGC statistics are provided in **Supplementary Table 3**.

BGCs were identified in MAGs retrieved from most phyla, namely 11 of 16 archaeal phyla and 52 out of 73 bacterial phyla (**Supplementary Table 3**). For bacteria **(Figure 2c)**, the most BGC-rich phyla are highly understudied^44, 48, 49^: the candidate phylum SZUA-182 (normalized counts of up to 12), Myxococcota (n ≈ 6) and DSWW01 (n ≈ 6). In agreement with their prevalence in cold seep sediments (**Figure 1b and Supplementary Figure 1**) and known capacity to produce specialized metabolites^7, 44, 48^, MAGs from phyla of Proteobacteria, Acidobacteriota, Planctomycetota, Desulfobacterota and Bacteroidota also encoded numerous biosynthetic gene clusters (**Figure 2c and Supplementary Table 3**). For archaea, EX4484-52 and Micrarchaeota from the DPANN superphylum (n ≈ 2) were the most BGC-rich **(Figure 2d)**, in agreement with other observations^44^. Thermoproteota, Halobacteriota, Asgardarchaeota, Thermoplasmatota, Altarchaeota, Hadarchaeota, Nanoarchaeota, PWEA01, and QMZS01 were also inferred to encode RiPPs, Terpenes, NRPS, polyketide synthases and other NPs.

The 2,865 BGCs clustered into 296 gene cluster families (GCFs) and 1,990 singletons **(Figure 2e)** based on BiG-SCAPE groupings^46^. Only three of the 2,865 cold seep BGCs grouped into GCFs together with reference BGCs^50^, specifically two NRPS (GCF_00393 and GCF_02819) and Other PKSs (GCF_01844). Most BGCs have unique structures and fewer similarity region modules (5-25%) compared to reference BGCs, and more are individually dispersed in the network (e.g. the 19 BGCs from PKS-NRPS hybrids; **Supplementary Figure 2**). We further compared the cold seep sediment BGCs with those from 16 other habitats (**Supplementary Figure 3**), including three artificial environments (bioreactors and wastewater), six host-associated environments (e.g. rumen and human gut), and seven natural environments (e.g. soil, freshwater and seawater)^24, 34, 41, 43, 51^. These data showed the relative proportions and dominant types of BGCs differ in cold seep sediments from other environments (**Supplementary Figure 3**), further highlighting the uniqueness of the deep biosphere.

### Natural products likely influence community dynamics and biogeochemical processes in cold seeps

We analyzed the potential function of the NPs based on wider integration with the literature. A large proportion of the BGCs likely encode for antimicrobial compounds, serving as chemical weapons for host defense and competition within the microbial community. Indeed, the most frequently predicted RiPPs were bacteriocins **(Figure 3a and Supplementary Table 3)**, which bacteria typically produce to inhibit and thereby outcompete other, often closely related, bacteria^52^. Also abundant were BGCs for various types of peptides, such as lasso peptides, thiopeptides, and lanthipeptides, which often have antimicrobial activities^53^. Among them, thiopeptides, also known as thiazolyl peptides, are structurally complex natural products with exquisite antibacterial activities and ability to overcome antibiotic resistance^54^. For the class of Others **(Figure 3a)**, aryl polyenes and beta-lactones were at high prevalence, indicating cold seep microbes synthesize NPs both for antimicrobial warfare and wider purposes such as protection against environmental stresses^37^. For example, exposure of cells to oxidative and reductive stress in high hydrostatic pressure cold seep environments can lead to the production of excess ROS, potentially causing cellular damage^55, 56^. The antioxidant action of aryl polyenes (APEs, like carotenoids) in microbes can scavenge external ROS, thereby preventing more extensive damage to essential cellular molecules and increasing bacterial fitness^19, 57^.

**Figure 3.**
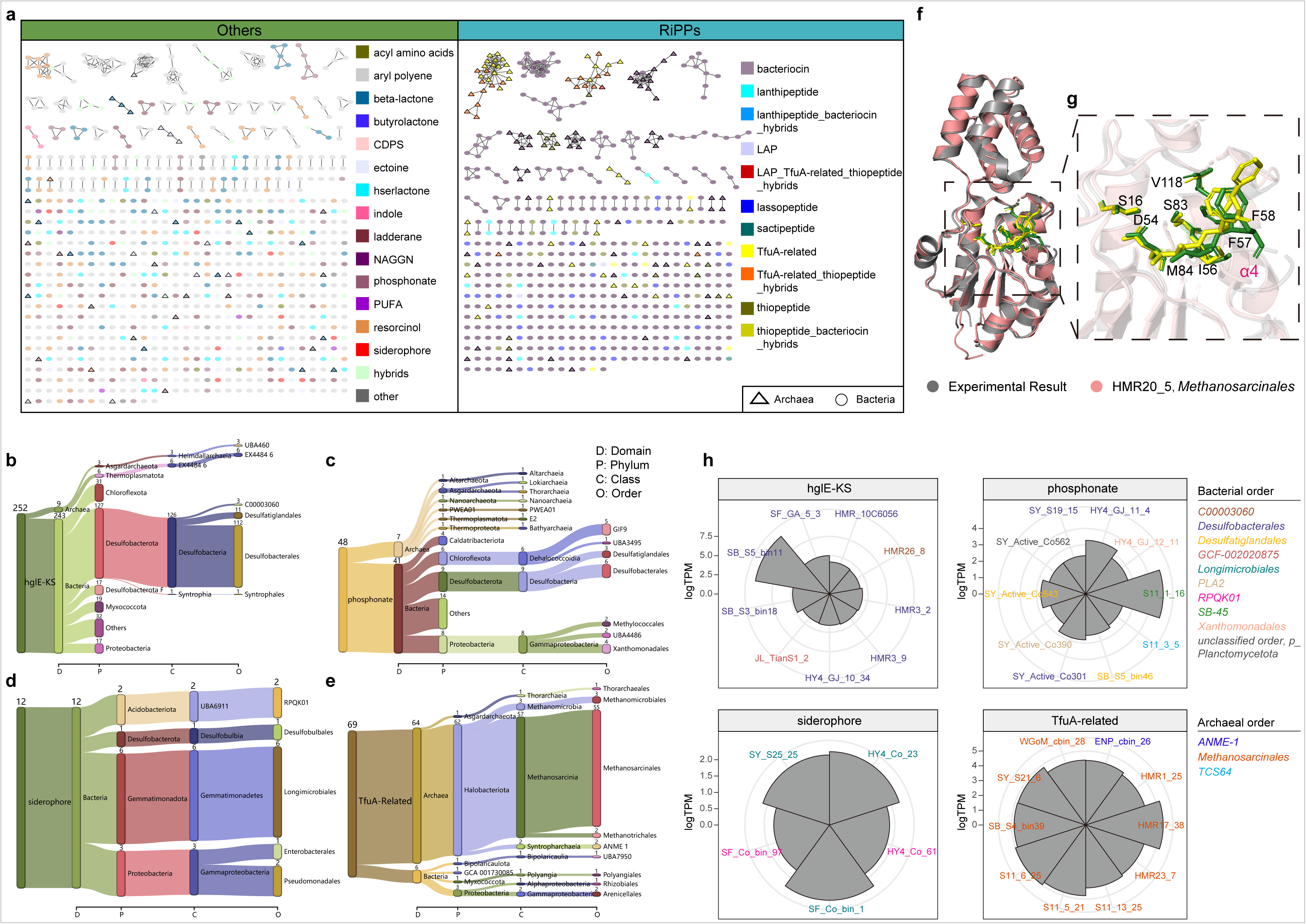
Biosynthetic gene clusters potentially influencing community dynamics and biogeochemical processes in cold seeps. **(a)** Detailed diversity of Others and RiPPs in archaeal and bacterial MAGs based on similarity networks. Nodes are colored by biosynthetic class. BGC distributions across different taxonomic levels, including (**b)** heterocyst glycolipid synthase-like PKS (hglE-KS), (**c)** phosphonate, (**d)** siderophore, and (**e)** TfuA-related. **(f-g)** Structural comparison between a predicted TfuA-related McrA-glycine thioamidation proteins and the experimentally determined structure (6XPB, the crystal structure of TfuA involved in peptide backbone thioamidation from *Methanosarcina acetivorans*). **(f)** The overall structural comparision showing a unique di-domain fold. The presumptive active site residues of experimental and predicted structure are shown as green and yellow sticks, respectively. **(g)** Close-up view of the putative active site of TfuA-related. Residues predicted to form the ThiS-binding pocket, and the α-helix that is implicated to mediate interactions with YcaO is marked as α4. **(h)** Wind rose diagram showing expression levels of BGCs from the top 10 or 5 abundant microbes related to N, P, Fe, and CH_4_ cycling. Transcript abundances are represented in the units of transcripts per million (TPM).

The NPs are also likely to be critical for the capacity of cold seep microbes to acquire nutrients and mediate biogeochemical cycling. For example, most diazotrophs with the phylum Desulfobacterota^58, 59^ (Desulfobacterales, Desulfatiglandales, C00003060 or SEEP-SRB1c and Syntrophales, **Figure 3b**) encoded heterocyst glycolipid synthase-like PKS (HglE-KS), which contributes to the synthesis of heterocyst-specific glycolipids (Hgls) to protect nitrogenase from oxygen damage^60, 61^. Six archaeal and 13 bacterial phyla also encoded BGCs to produce phosphonates **(Figure 3c)**, which have diverse structural, antimicrobial, and storage roles and are central to marine phosphorus cycling^62^. Certain taxa, including multiple Gemmatimonadota MAGs **(Figure 3d)**, encode siderophores to scavenge the presumably limiting micronutrient iron^63^.

Also notable are the BGCs encoded by Halobacteriota, the main players of anaerobic oxidation of methane occurring in cold seeps^7^. A total of 62 TfuA-related BGCs **(Figure 3e)** were encoded by the orders Methanosarcinales, Methanomicrobiales, Methanotrichales, and ANME-1 from this phylum. These gene clusters encode TfuA-like and YcaO proteins^64^ that mediate the post-translational modification of methyl-coenzyme M reductase (MCR)^65, 66^, a large protein ubiquitous to methanogenic and methanotrophic archaea that catalyzes the anaerobic production and consumption of methane^66^. Indeed, the predicted amino acid sequences of the TfuA-like protein is related to the McrA-glycine thioamidation proteins **(Supplementary Figure 4).**

Structural predictions confirmed that these cold seep proteins are identical in secondary and tertiary structure to experimentally determined thioamidation proteins^65^ and form the required substrate-binding pockets **(Figure 3f**, **3g, and Supplementary Figure 5)**. Thus, we predict that cold seep Halobacteriota encode YcaO and TfuA proteins to help catalyse the thioamidation of the MCR, thereby enabling anaerobic methane oxidation.

### BGCs from key players in cold seeps are widely distributed, highly expressed, and depth stratified

At the ecosystem level, the BGCs from the 63 different phyla were encoded in at least one cold seep site, though their abundance and occupancy varied **(Figure 4a, Supplementary Figure 6 and Supplementary Table 4)**. Gene-centric surveys suggested that the phosphonate biosynthesis gene from Caldatribacteriota (ENP_sbin_8_3) was the most abundant across different samples. Notably, the abundance of BGCs in Asgardarchaeota, Hadarchaeota, Halobacteriota, Thermoproteota, Actinobacteriota, Aerophobota, Caldatribacteriota, Desulfobacterota, and UBA6262 were also globally high across different sites, with the mean abundance greater than 500 GPM **(Figure 4b; see Methods)**. For archaea, a high abundance of BGCs from Halobacteriota, Asgardarchaeota, Thermoproteota, and Thermoplasmatota were observed across globally distributed cold seep sediments in almost every depth.

**Figure 4.**
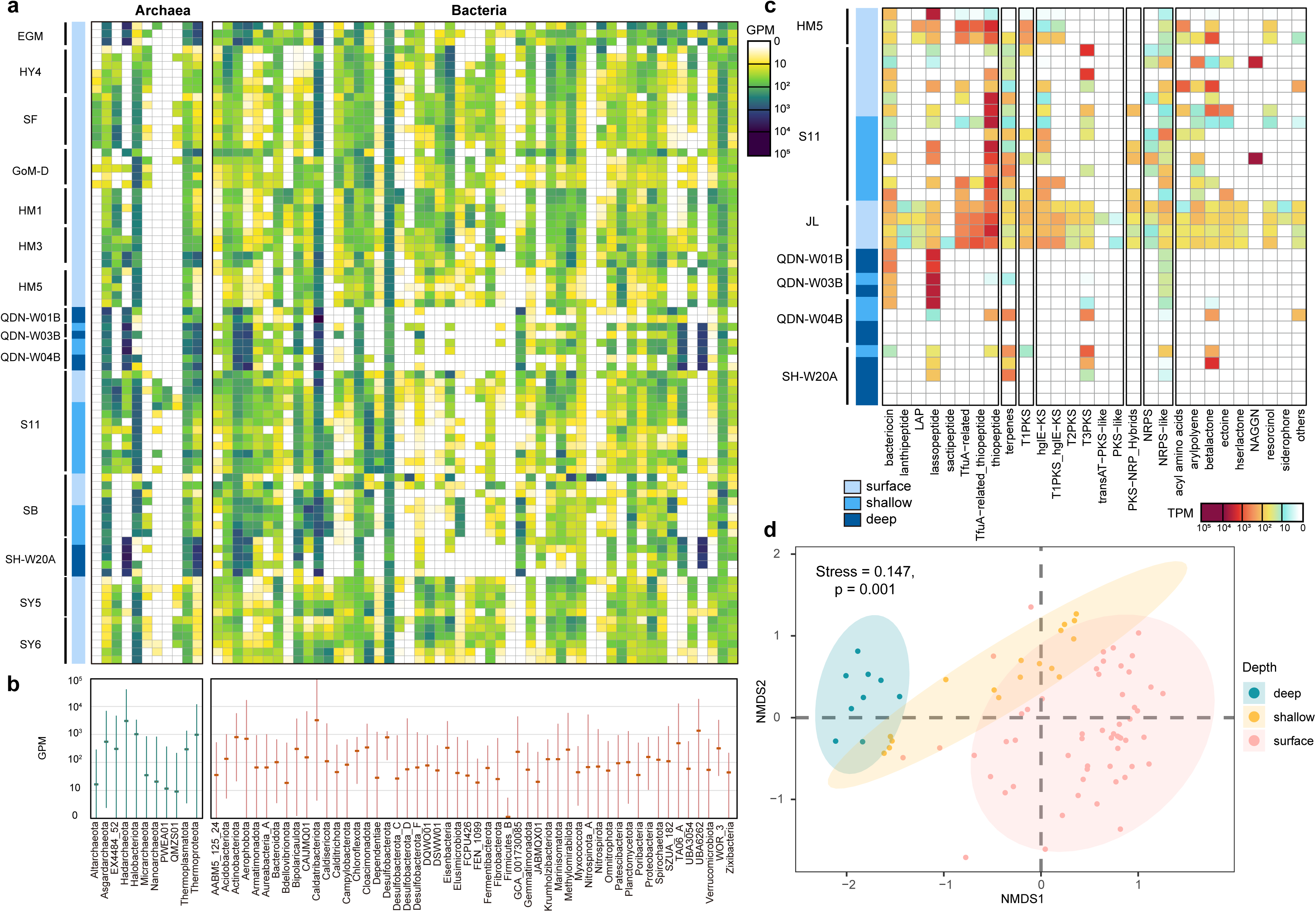
Abundance and expression of biosynthetic gene clusters across different phyla and different BGC types in the cold seep sediments. **(a)** The heatmap shows the average abundance of BGCs for each phylum. Sediment depth is grouped into: surface, <1 mbsf; shallow, 1–10 mbsf; deep, >10 mbsf. **(b)** Mean relative abundances of BGCs for each phylum. The bars indicate the minima and maxima of BGC abundances. BGC abundances are represented in the units of genes per million (GPM). **(c)** The heatmap shows the average transcript abundance of BGCs for each type. Transcript abundances are represented in the units of transcripts per million (TPM). **(d)** NMDS analysis of a Bray-Curtis dissimilarity matrix calculated from BGC abundances. ANOSIM was applied to test BGC differences in microbial communities among different sediment depths (surface, <1 mbsf; shallow, 1–10 mbsf; deep, >10 mbsf), using a 999-permutation test. Detailed data are in **Supplementary Tables 4-5**.

Metatranscriptome analyses confirmed expression of BGCs from various microorganisms at different sites and depths (**Figure 4c, Supplementary Figure 7 and Supplementary Table 5**). For example, 42 bacterial (>80%) and 7 archaeal phyla (>63%) expressed BGCs in the Jiaolong cold seep sediments. In bacteria, BGC expression was dominated by Actinobacteriota, Bipolaricaulota, Chloroflexota, Desulfobacterota, and WOR-3 **(Supplementary Figure 7)**, especially biosynthesis genes for bacteriocins, beta-lactones, aryl polyenes, and polyketides with inferred antibiotic and oxidative stress resistance activities^39^. For example, these genes were highly expressed in Haima cold seep sediments (HM5 and S11), especially those for T3PKS (up to 932,163 TPM**; see Methods**) and aryl polyene (up to 842,651 TPM) synthesis **(Figure 4c)**. The genes to produce nitrogenase-protecting glycopeptides (up to 171,389 TPM in Desulfobacterales), siderophores (up to 201 TPM in Longimicrobiales) and phosphonates (up to 54,264 TPM in SB-45) were also expressed (**Figure 3h**). Among archaea, we also observed a high expression of various Halobacteriota BGCs, including thiopeptides associated with antimicrobial defenses (up to 502,455 TPM in ANME-1) and TfuA-related genes for methyl-CoM reductase biosynthesis (up to 172,569 TPM in Methanosarcinales).

To determine the distribution characteristics of biosynthetic gene clusters, each metagenome was categorized in terms of its sediment depth (i.e. surface, <1 mbsf; shallow, 1–10 mbsf; deep, >10 mbsf). BGC compositions were stratified by depth **(Figure 4d)**, with significantly differences being observed among surface, shallow and deep layers (*p* LJ=LJ 0.001, R LJ=LJ 0.459, 999-permutations test). Likewise, we observed depth-related differences in BGC expression **(Supplementary Figure 7**). In general, very few BGC transcripts were detected at deeper sediment sites, though some transcripts were still highly expressed such as a lasso peptide biosynthesis gene (up to 482,852 TPM) at QDN W01B and W03B sites.

### Sediment metabolomes confirm BGC expression and NP novelty

To generate chemical evidence that confirms the presence of diverse secondary metabolites, we utilized untargeted mass-spectrometry-based metabolomics to examine metabolomes in the sediment samples of Qiongdongnan cold seeps (QDN; **Figure 1**) at depths of 0-30 cmbsf. This resulted in the identification of 9,145 lipophilic and 1,061 hydrophilic molecular features based on MS/MS spectra comparisons with the GNPS online platform^67^ **(Figure 5a, Supplementary Tables 6 and 7)**. A total of 3,054 molecular features (29.9%) were shared between more than two depths. Most other metabolites were only detected in sediment at a specific depth (**Supplementary** Figures 8-9), suggesting spatial stratification of cold seep sediment metabolomes. Like BGCs detected in the metagenomes, the generated molecular networks **(Figure 5b and Supplementary Figure 8)** displayed a significant number of singleton nodes (5145, 50.4%), suggesting that a substantial amount of structurally dissimilar compounds were detected.

**Figure 5.**
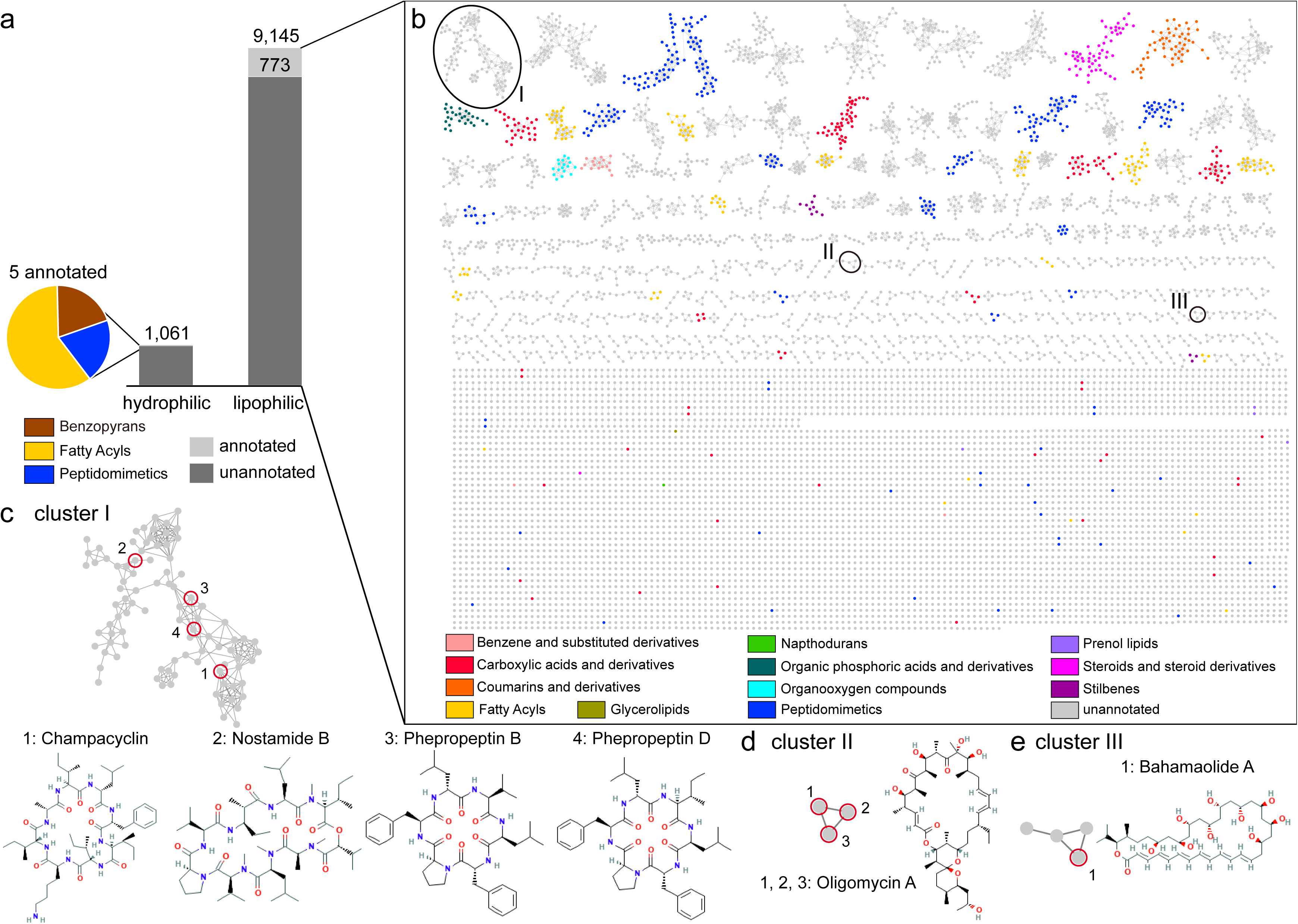
Metabolomes in Qiongdongnan cold seep sediments. **(a)** Annotated and unannotated molecular features were classified into two groups: hydrophilic and lipophilic metabolites. Annotated hydrophilic metabolites are grouped into three classes. **(b)** A molecular network of 9,145 nodes was created, 773 of which fall into 12 molecular families and are indicated with different colors while the remaining were colored in grey to represent unannotated lipophilic metabolites. **(c-e)** Clusters of notable identified lipophilic metabolites. Cluster I **(c)** were peptides, with nodes identified to cyclic peptides. The cluster II (d) and III **(e)** belonged to macrolides. Detailed data are in **Supplementary Tables 6-9**.

Despite extensive manual inspection, different database searches, and other cheminformatic predictions, most detected molecular features in QDN sites could not be matched with known compounds^68–70^. A total of 778 molecular features **(Figure 5)** were classified into thirteen classes by the initial structural predictions based on MS/MS spectra comparisons with chemical structure databases (including HMDB, SUPNAT, CHEBI, DRUGBANK, and FooDB) by the MolNetEnhancer workflow^70^. This suggests that most molecular features remain unidentified and there is a vast, unexplored chemical space associated with sediment metabolomes in cold seeps. The 773 annotated lipophilic molecular features were classified into twelve classes **(Figure 5b and Supplementary Table 7)**, with 436 being identified as peptides, including 356 cyclic peptides (cyclopeptides) and 80 oligopeptides.

A total of 68 hydrophilic and 462 lipophilic compounds were predicted with the database Natural Products Atlas by SNAP-MS^71^ **(Supplementary Tables 8 and 9)**. A range of peptides were predicted in the lipophilic network **(Figure 5c and Supplementary Table 10)**, including cyclic hexapeptides and octapeptides, the antimicrobial compound champacyclin^72^, and the proteasome inhibitors phepropeptin B and D^73, 74^ (**Figure 5c**). Nostamides (**Figure 5c and Supplementary Figure 10**), predicted to be synthesised by 16 NRPS biosynthetic gene clusters from five phyla (Calditrichota, Chloroflexota, Cloacimonadota, Planctomycetota and Proteobacteria), were also identified. These cyclic hexapeptides, also known as anabaenopeptins, can be potent inhibitors of blood clot stabilizing carboxypeptidases^75^. Besides, there were other classes of bioactive natural products **(Figure 5d and 5e)**, such as macrolides, bahamaolide A with antifungal activity ^76^ and oligomycin A with cytotoxicity^77^.

## Conclusions

This meta-omic study, bridging metagenomic, metatranscriptomic, and metabolomic analysis, suggests a vast diversity of natural products are produced in an underexplored deep-sea chemosynthetic ecosystem. BGCs were encoded by 63 microbial phyla from nine seeps, with 7 archaeal and 42 bacterial phyla expressing these genes in at least one sampling location. In turn, this expands our knowledge of the biosynthetic potential of uncultivated anaerobic microbes, especially the rarely investigated archaeal domain^39, 78, 79^. The derived natural products likely influence community dynamics, for example through their inferred roles in antimicrobial warfare, oxidative stress defences, and potentially biofilm formation, but also likely also influence biogeochemical cycling for example through their roles in facilitating methyl-CoM reductase biosynthesis in anaerobic methanotrophic archaea. Together, the metagenomic and metabolomic analysis suggest these microorganisms synthesize many unknown metabolites, though it is challenging with current understanding to accurately predict what NPs are synthesized from BGCs. Further biochemical analysis, for example through heterologous expression of BGCs and production of NPs, will be required to gain a more comprehensive understanding of these links and explore the pharmaceutical potential of the cold seep metabolome.

## Methods

### Collection of metagenomes, metatranscriptomes and MAGs

For this study, the 81 metagenomes and 33 metatranscriptomes used were derived from nine globally distributed cold seeps **(Figure 1a and in Supplementary Table 1)**. These sites are as follows: Eastern Gulf of Mexico (EGM), Northwestern Gulf of Mexico (GOM), Scotian Basin (SB), Haima seep (HM1, HM3, HM5, S11, SY5, SY6), Haiyang4 (HY4), Site F cold seep (SF), Qiongdongnan Basin (QDN), Shenhu area (SH), and Jiaolong cold seep (JL). Detailed descriptions of the sampling sites and sequencing information for the metagenomic and metatranscriptomic data can be found in our previous work^7, 45, 58, 80^. Paired-end raw reads were quality-controlled by trimming primers and adapters and filtering out artifacts and low-quality reads using the Read_QC module within the metaWRAP pipeline (v1.3.2; –skip-bmtagger)^81^. Clean reads from each sample at individual depths were assembled, and clean reads from each sampling station from all depths were co-assembled, both using MEGAHIT (v1.1.3; default parameters, length > 1000 bp). For each assembly, contigs were binned using the binning module (parameters: –maxbin2 –concoct –metabat2) and consolidated into a final bin set using the Bin_refinement module (parameters: –c 50 –x 10) within metaWRAP. The quality of the obtained MAGs was estimated by the lineage-specific workflow of CheckM (v1.0.12)^82^. MAGs estimated to be at least 50% complete and with less than 10% contamination were retained. MAGs reconstructed from individual assemblies and co-assemblies were combined and dereplicated for strains-level clustering using dRep (v3.2.2)^83^ with an average nucleotide identity (ANI) cutoff value of 99%, resulting in 2,479 strains-level representative MAGs. These MAGs obtained were taxonomically annotated using the GTDB-Tk (v1.5.0)^84^ against Genome Taxonomy Database GTDB (release 06-RS202). Phylogenetic trees produced by GTDB-Tk were uploaded to iTOL (v6.7.5; https://itol.embl.de/)^85^ for visualization.

### BGCs mining and net-work analysis

BGCs were identified from MAGs using antiSMASH (v5.1.2)^27^ using the following parameters: -fullhmmer -cb-general -cb-subclusters -cb-knownclusters -genefinding-tool prodigal-m -clusterhmmer -asf -smcog-trees -pfam2go. Only BGCs on contigs of > 10 kb were considered. BiG-SCAPE (v1.1.0)^46^ was run in -auto mode with -mibig enabled to identify BGCs families. Networks using the similarity threshold of 0.3 were examined, generating proposed BGC families (Gene Cluster Family, GCF).

### Phylogenetic analyses of functional genes

The TfuA-like phylogenetic tree was produced by aligning the predicted TfuA-like protein sequences with 25 reference sequences. All sequences were aligned using MAFFT (v7.505, –auto option)^86^ and gap sequences were trimmed using TrimAl (v1.2.59, –gappyout option)^87^. The maximum likelihood tree was constructed with IQ-Tree (v2.1.2; with best-fit models and 1000 bootstrapped replicates)^88^ on the CIPRES server^89^. Three-dimensional structures of TfuA-like protein sequences were predicted using ColabFold by combining the fast homology search of MMseqs2 with AlphaFold2 ^90, 91^.

### BGCs abundance calculations and in situ activities

The relative abundance of BGCs across difference metagenomes were calculated from the combined BGC catalog (n = 2,865) with the program Salmon (v1.8.0)^62^ in the mapping-based mode (parameters: -validateMappings -meta). GPM (genes per million) values were used as a proxy for BGC abundance. The ribosomal RNAs in quality-filtered metatranscriptomic reads were removed by comparing with rRNA sequences in the Rfam and Silva databases using SortMeRNA (v4.2.0)^92^. Preprocessed reads were mapped to the combined BGC catalog to generate read count quantification TPM (Transcripts Per Million) of each transcript using Salmon (v1.8.0)^93^. The sum of all TPM values is the same in all samples and gene expression levels between samples are comparable in principle, but it is noted that the differences in the abundance of some highly expressed genes are likely to affect the relative abundance of all transcripts in a sample, thus may leading to high TPM values of some genes in a sample^94^.

### Analysis of metabolites from sediment samples

*Extraction of metabolites from sediment samples.* Metabolite extraction was conducted using established protocols^68^. Hydrophilic and lipophilic metabolites were prepared separately. About two grams of each sediment sample stored at −80 °C were taken and directly suspended in 10 ml of 1:1 dichloromethane/methanol (vol/vol) in a glass tube. The tubes were shaken carefully and sonicated for 10 min. The supernatant was collected. Fresh solvents were added to the tubes twice to repeat the extraction process and all supernatants were combined. The mixtures were filtrated using 0.45 μm filters and solvents were then removed by rotary evaporation (30 °C). The obtained crude extracts were redissolved by adding 20 ml of chloroform/methanol/water (2:1:1, vol/vol/vol). The mixtures were vortex-mixed for 2 min and centrifuged at 10,000 g for 10 min at 4 °C. Both the upper (water-soluble metabolites) and the bottom phases (lipid-soluble metabolites) were recovered and vacuum-dried using a vacuum evaporator. The hydrophilic and the lipophilic metabolites were reconstituted using 2 ml of acetonitrile/water (1:1, vol/vol) and isopropanol/acetonitrile/water 4:3:1 (vol/vol/vol), respectively. Two reagent blank samples (one hydrophilic and one lipophilic sample) were also prepared using the same procedure while no sediment sample was added in the first step.

#### Analysis of metabolites samples using LC-MS/MS

Liquid chromatography tandem-mass spectrometry (LC-MS/MS) analyses were performed on a Vion^TM^ ion-mobility quadrupole time-of-flight mass spectrometer (Waters, MA, USA) with an electrospray interface and coupled with an Acquity UPLC system (Waters, MA, USA). The ESI conditions were as follows: capillary voltage 2500 V, source temperature 120 °C, desolvation temperature 350 °C, cone gas flow 80 L/h, desolvation gas flow of 850 L/h. Detection was performed in positive ion mode in the m/z range of 50–2000 with a scan time of 0.2 s. The MS^E^ acquisition mode, which allows simultaneous acquisition of full scan data (low-energy scan, 6 eV) and collision-induced fragmentation data (high-energy scan, ramp from 25 to 35 eV), was used. Leucine-enkephalin (Sigma–Aldrich, Steinheim, Germany) of 200 ng/mL was used as a reference and was introduced into the system at a flow rate of 10 µl/min. The separation of hydrophilic metabolites was carried out on an ACQUITY UPLC BEH Amide column (2.1 × 150 mm, particle size 1.7 µm). Acetonitrile and water (both with 0.1% (vol/vol) formic acid) were used as mobile phases A and B, respectively. The column temperature was set to 30 °C, the flow rate to 0.4 ml/min, the injection volume to 3 µl, and the autosampler temperature to 10 °C. The following elution gradient was used: 0-6 min, 1 % to 60 % B; 6-8 min, 60 % to 1 % B; and 1% B was kept for 2 min. The gradient was run four times without injecting any sample to equilibrate the column. All samples were diluted ten times before analysis. Lipophilic metabolites were separated on an ACQUITY UPLC CSH C18 column (2.1 × 100 mm, particle size 1.7 µm). Acetonitrile/water (60:40, vol/vol) with 10 mM ammonium formate and 0.1% (vol/vol) formic acid was used as mobile phase A. Isopropanol/acetonitrile (90:10, vol/vol) with 10 mM ammonium formate and 0.1% (vol/vol) formic acid was used as mobile phase B. The column temperature was set at 50 °C, the flow rate at 0.4 ml/min, the injection volume at 3 µl, and the autosampler temperature at 10 °C. The following elution gradient was used: 0-2 min, 40 % to 43 % B; 2–2.1 min, 43 % to 50 % B; 2.1–10 min, 50 % to 54 % B; 10–10.1 min, 54 % to 70 % B; 10.1–14 min, 70 % to 99 % B; 14–14.1 min, 99 % to 40 % B; and 14.1–16 min, kept 40% B. The gradient was run four times without injecting any sample to equilibrate the column.

#### Metabolite annotation using GNPS platform

Raw data generated from LC-MS/MS analysis were converted into the mgf format using UNIFI v. 1.8 (Waters). The mgf files were uploaded and deposited in the MassIVE Public GNPS (Global Natural Products Social Molecular Networking) data set (https://massive.ucsd.edu/)^67, 69^. Firstly, the MS/MS data were analyzed with the GNPS pipeline (METABOLOMICS-SNETS) using the following parameters: precursor ion mass tolerance = 0.01 Da, fragment ion mass tolerance = 0.01 Da, minimal pairs cosine = 0.7, minimum matched fragment ions = 6, and minimum cluster size = 1. The options “Search Analogs” was checked. One no-injection blank sample and one reagent blank sample were put in G6 to filter the contaminants and/or noises from UPLC analysis as well as sample preparation procedures. Compounds were identified using the GNPS reference library, which contains over 220,000 curated spectra. Library matches were assigned if they shared at least 6 MS/MS peaks and had a similarity score above 0.7. For all spectral matches, the generated mirror spectra plots were manually investigated. Secondly, the Dereplicator tool^95^ was used for the identification of known peptidic natural products from the generated molecular network. The precursor ion mass tolerance and fragment ion mass tolerance were both set as 0.02 Da. The option “Search Analogs (VarQuest)” was checked. The molecular networks were also subjected to MS2LDA (http://ms2lda.org/) workflow^96, 97^. The bin width was set as 0.01. The minimum MS2 intensity and LDA free motifs were set as 100 and 200, respectively. Subsequently, the Network Annotation Propagation (NAP) workflow was used^98^. The following parameters were applied: 10 first candidates for consensus score, accuracy for exact mass candidate search within 15 ppm, and search against structure databases including HMDB, SUPNAT, CHEBI, DRUGBANK, and FooDB. Lastly, the results generated from the above-mentioned workflows were integrated by MolNetEnhancer into the generated molecular network^70^. All the nodes annotated by the library search or in silico prediction were submitted to chemical classification using ClassyFire hierarchical chemical ontology software^99^. The molecular network was then visualized using Cytoscape (v3.8.2)^100^. Compound family annotations were also carried out through matching chemical similarity grouping in the Natural Products Atlas to grouping of mass spectrometry features from molecular networks by Structural similarity Network Annotation Platform for Mass Spectrometry (SNAP-MS)^71^.

### Statistical analyses

All statistical analyses were performed in R (v4.1.3). Beta diversity of biosynthetic gene clusters was calculated using vegan package (v2.5–6)^101^. Shapiro–Wilk and Bartlett’s tests were employed to test data normality and homoscedasticity prior to other statistical analysis. Non-metric multidimensional scaling (NMDS) was used to reduce dimensionality using the function capscale, based on Bray–Curtis dissimilarities generated with gene abundances (GPM) values using the vegdist function. The groupings of cold seep sediments into three different sample depths (surface: <1 mbsf, shallow: 1-10 mbsf, and deep: >10 mbsf) were individually verified using Analysis of Similarity (ANOSIM), performed with 999 permutations based on Bray–Curtis dissimilarity.

### Data availability

MAGs, BGCs and other related information have been uploaded to figshare (10.6084/m9.figshare.23364041). Raw data generated from LC-MS/MS analysis were converted into the mgf format using UNIFI v. 1.8 (Waters). The mgf files were uploaded and deposited in the MassIVE Public GNPS (Global Natural Products Social Molecular Networking) data set (https://massive.ucsd.edu/). The accession numbers for polar and nonpolar metabolites data are MSV000090350 and MSV000090349, respectively.

### Code availability

The present study did not generate codes, and mentioned tools used for the data analysis were applied with default parameters unless specified otherwise.

## Acknowledgments

The work was supported by the Natural Science Foundation of Fujian Province (No. 2023J06042), Scientific Research Foundation of Third Institute of Oceanography, MNR (No. 2022025 and No. 2023022), State Key Laboratory of Marine Geology, Tongji University (No. MGK202303), and China Postdoctoral Science Foundation (2023M734096). We thank Chuwen Zhang, Chuan Huang, Max Cryle and Lu Zhang for providing valuable comments.

## Author contributions

X.D. designed this study. X.D., T.Z., Y.P., and X.L. performed metagenomic and metatranscriptomic analyses. T.Z., W.W., X.C., J.X., and Z.S contributed to sediment metabolomes. X.D., T.Z., W.W., Y.H., Z.G., J.X., Z.S and C.G. interpreted the data. T.Z., X.D., and C.G. wrote the paper, with input from other authors.

## References

1. Boetius, A. & Wenzhöfer, F. Seafloor oxygen consumption fuelled by methane from cold seeps. Nature Geoscience 6, 725–734 (2013).

2. Peckmann, J. & Thiel, V. Carbon cycling at ancient methane-seeps. Chemical Geology 205, 443–467 (2004).

3. Knittel, K. & Boetius, A. Anaerobic oxidation of methane: progress with an unknown process. Annu Rev Microbiol 63, 311–334 (2009).

4. Gibson, R. et al. Ecology of cold seep sediments: Interactions of fauna with flow, chemistry and microbes. An Annual Review 43, 1–46 (2005).

5. Emil Ruff, S. in Marine Hydrocarbon Seeps. (eds. A. Teske & V. Carvalho) 1–19 (Springer International Publishing, Cham; 2020).

6. Ruff, S.E. et al. Global dispersion and local diversification of the methane seep microbiome. Proc Natl Acad Sci U S A 112, 4015–4020 (2015).

7. Dong, X. et al. Evolutionary ecology of microbial populations inhabiting deep sea sediments associated with cold seeps. Nat Commun 14, 1127 (2023).

8. Peng, Y. et al. Viruses in deep-sea cold seep sediments harbor diverse survival mechanisms and remain genetically conserved within species. bioRxiv, 2023.2003.2012.532262 (2023).

9. Anderson, R.E. Tracking Microbial Evolution in the Subseafloor Biosphere. mSystems 6, e0073121 (2021).

10. Newman, D.J. & Cragg, G.M. Natural Products as Sources of New Drugs over the Nearly Four Decades from 01/1981 to 09/2019. J Nat Prod 83, 770–803 (2020).

11. Scherlach, K. & Hertweck, C. Chemical Mediators at the Bacterial-Fungal Interface. Annu Rev Microbiol 74, 267–290 (2020).

12. Mullis, M.M., Rambo, I.M., Baker, B.J. & Reese, B.K. Diversity, Ecology, and Prevalence of Antimicrobials in Nature. Front Microbiol 10, 2518 (2019).

13. Abrudan, M.I. et al. Socially mediated induction and suppression of antibiosis during bacterial coexistence. Proc Natl Acad Sci U S A 112, 11054–11059 (2015).

14. van Bergeijk, D.A., Terlouw, B.R., Medema, M.H. & van Wezel, G.P. Ecology and genomics of Actinobacteria: new concepts for natural product discovery. Nat Rev Microbiol 18, 546–558 (2020).

15. Wright, E.S. & Vetsigian, K.H. Inhibitory interactions promote frequent bistability among competing bacteria. Nat Commun 7, 11274 (2016).

16. Xia, L. et al. Biosynthetic gene cluster profiling predicts the positive association between antagonism and phylogeny in Bacillus. Nat Commun 13, 1023 (2022).

17. Cornforth, D.M. & Foster, K.R. Antibiotics and the art of bacterial war. Proc Natl Acad Sci U S A 112, 10827–10828 (2015).

18. Neilands, J.B. Siderophores. Arch Biochem Biophys 302, 1–3 (1993).

19. Johnston, I. et al. Identification of essential genes for Escherichia coli aryl polyene biosynthesis and function in biofilm formation. NPJ Biofilms Microbiomes 7, 56 (2021).

20. Sadeghi, A. et al. Diversity of the ectoines biosynthesis genes in the salt tolerant Streptomyces and evidence for inductive effect of ectoines on their accumulation. Microbiol Res 169, 699–708 (2014).

21. Woon, S.A. & Fisher, D. Antimicrobial agents - optimising the ecological balance. BMC Med 14, 114 (2016).

22. Junkins, E.N., McWhirter, J.B., McCall, L.-I. & Stevenson, B.S. Environmental structure impacts microbial composition and secondary metabolism. ISME Communications 2, 15 (2022).

23. Shaffer, J.P. et al. Standardized multi-omics of Earth’s microbiomes reveals microbial and metabolite diversity. Nat Microbiol 7, 2128–2150 (2022).

24. Crits-Christoph, A., Diamond, S., Butterfield, C.N., Thomas, B.C. & Banfield, J.F. Novel soil bacteria possess diverse genes for secondary metabolite biosynthesis. Nature 558, 440–444 (2018).

25. Doroghazi, J.R. et al. A roadmap for natural product discovery based on large-scale genomics and metabolomics. Nat Chem Biol 10, 963–968 (2014).

26. Cimermancic, P. et al. Insights into secondary metabolism from a global analysis of prokaryotic biosynthetic gene clusters. Cell 158, 412–421 (2014).

27. Blin, K. et al. antiSMASH 5.0: updates to the secondary metabolite genome mining pipeline. Nucleic Acids Res 47, W81–W87 (2019).

28. Hannigan, G.D. et al. A deep learning genome-mining strategy for biosynthetic gene cluster prediction. Nucleic Acids Res 47, e110 (2019).

29. Sanchez, S. et al. Expansion of novel biosynthetic gene clusters from diverse environments using SanntiS. bioRxiv, 2023.2005.2023.540769 (2023).

30. Carroll, L.M. et al. Accurate *de novo* identification of biosynthetic gene clusters with GECCO. bioRxiv, 2021.2005.2003.442509 (2021).

31. Medema, M.H., de Rond, T. & Moore, B.S. Mining genomes to illuminate the specialized chemistry of life. Nat Rev Genet 22, 553–571 (2021).

32. Salamzade, R. et al. Evolutionary investigations of the biosynthetic diversity in the skin microbiome using lsaBGC. Microb Genom 9 (2023).

33. Donia, M.S. et al. A systematic analysis of biosynthetic gene clusters in the human microbiome reveals a common family of antibiotics. Cell 158, 1402–1414 (2014).

34. Anderson, C.L. & Fernando, S.C. Insights into rumen microbial biosynthetic gene cluster diversity through genome-resolved metagenomics. Commun Biol 4, 818 (2021).

35. Mendes, R. et al. Deciphering the rhizosphere microbiome for disease-suppressive bacteria. Science 332, 1097–1100 (2011).

36. Rust, M. et al. A multiproducer microbiome generates chemical diversity in the marine sponge Mycale hentscheli. Proc Natl Acad Sci U S A 117, 9508–9518 (2020).

37. Paoli, L. et al. Biosynthetic potential of the global ocean microbiome. Nature 607, 111–118 (2022).

38. Schorn, M.A. et al. Sequencing rare marine actinomycete genomes reveals high density of unique natural product biosynthetic gene clusters. Microbiology (Reading) 162, 2075–2086 (2016).

39. Geller-McGrath, D. et al. Diverse secondary metabolites are expressed in particle-associated and free-living microorganisms of the permanently anoxic Cariaco Basin. Nat Commun 14, 656 (2023).

40. Tuttle, R.N. et al. Detection of Natural Products and Their Producers in Ocean Sediments. Appl Environ Microbiol 85 (2019).

41. Waschulin, V. et al. Biosynthetic potential of uncultured Antarctic soil bacteria revealed through long-read metagenomic sequencing. ISME J 16, 101–111 (2022).

42. Liu, Y. et al. A genome and gene catalog of glacier microbiomes. Nat Biotechnol 40, 1341–1348 (2022).

43. Nayfach, S. et al. A genomic catalog of Earth’s microbiomes. Nat Biotechnol 39, 499–509 (2021).

44. Galal, A. et al. A survey of the biosynthetic potential and specialized metabolites of archaea and understudied bacteria. Current Research in Biotechnology 5, 100117 (2023).

45. Han, Y. et al. A comprehensive catalog with 100 million genes and 3,000 metagenome-assembled genomes from global cold seep sediments. bioRxiv, 2023.2004.2010.536201 (2023).

46. Navarro-Munoz, J.C. et al. A computational framework to explore large-scale biosynthetic diversity. Nat Chem Biol 16, 60–68 (2020).

47. Wei, B. et al. Global analysis of the biosynthetic chemical space of marine prokaryotes. Microbiome 11, 144 (2023).

48. Gavriilidou, A. et al. Compendium of specialized metabolite biosynthetic diversity encoded in bacterial genomes. Nat Microbiol 7, 726–735 (2022).

49. Hoffmann, T. et al. Correlating chemical diversity with taxonomic distance for discovery of natural products in myxobacteria. Nat Commun 9, 803 (2018).

50. Medema, M.H. et al. Minimum Information about a Biosynthetic Gene cluster. Nat Chem Biol 11, 625–631 (2015).

51. Sharrar, A.M. et al. Bacterial Secondary Metabolite Biosynthetic Potential in Soil Varies with Phylum, Depth, and Vegetation Type. mBio 11 (2020).

52. Negash, A.W. & Tsehai, B.A. Current Applications of Bacteriocin. Int J Microbiol 2020, 4374891 (2020).

53. Shen, X., Mustafa, M., Chen, Y., Cao, Y. & Gao, J. Natural thiopeptides as a privileged scaffold for drug discovery and therapeutic development. Medicinal Chemistry Research 28, 1063–1098 (2019).

54. Vinogradov, A.A. & Suga, H. Introduction to Thiopeptides: Biological Activity, Biosynthesis, and Strategies for Functional Reprogramming. Cell Chem Biol 27, 1032–1051 (2020).

55. Poljsak, B., Suput, D. & Milisav, I. Achieving the balance between ROS and antioxidants: when to use the synthetic antioxidants. Oxid Med Cell Longev 2013, 956792 (2013).

56. Zhang, Y., Li, X., Bartlett, D.H. & Xiao, X. Current developments in marine microbiology: high-pressure biotechnology and the genetic engineering of piezophiles. Curr Opin Biotechnol 33, 157–164 (2015).

57. Schoner, T.A. et al. Aryl Polyenes, a Highly Abundant Class of Bacterial Natural Products, Are Functionally Related to Antioxidative Carotenoids. Chembiochem 17, 247–253 (2016).

58. Dong, X. et al. Phylogenetically and catabolically diverse diazotrophs reside in deep-sea cold seep sediments. Nat Commun 13, 4885 (2022).

59. Metcalfe, K.S., Murali, R., Mullin, S.W., Connon, S.A. & Orphan, V.J. Experimentally-validated correlation analysis reveals new anaerobic methane oxidation partnerships with consortium-level heterogeneity in diazotrophy. ISME J 15, 377–396 (2021).

60. Awai, K., Lechno-Yossef, S. & Wolk, C.P. in Lipids in Photosynthesis. (eds. H. Wada & N. Murata) 179–202 (Springer Netherlands, Dordrecht; 2009).

61. Garg, R. & Maldener, I. The Dual Role of the Glycolipid Envelope in Different Cell Types of the Multicellular Cyanobacterium Anabaena variabilis ATCC 29413. Front Microbiol 12, 645028 (2021).

62. Lockwood, S., Greening, C., Baltar, F. & Morales, S.E. Global and seasonal variation of marine phosphonate metabolism. ISME J 16, 2198–2212 (2022).

63. Kramer, J., Ozkaya, O. & Kummerli, R. Bacterial siderophores in community and host interactions. Nat Rev Microbiol 18, 152–163 (2020).

64. Malit, J.J.L., Wu, C., Liu, L.L. & Qian, P.Y. Global Genome Mining Reveals the Distribution of Diverse Thioamidated RiPP Biosynthesis Gene Clusters. Front Microbiol 12, 635389 (2021).

65. Liu, A. et al. Functional elucidation of TfuA in peptide backbone thioamidation. Nat Chem Biol 17, 585–592 (2021).

66. Nayak, D.D., Mahanta, N., Mitchell, D.A. & Metcalf, W.W. Post-translational thioamidation of methyl-coenzyme M reductase, a key enzyme in methanogenic and methanotrophic Archaea. Elife 6 (2017).

67. Nothias, L.F. et al. Feature-based molecular networking in the GNPS analysis environment. Nat Methods 17, 905–908 (2020).

68. Paglia, G. & Astarita, G. Metabolomics and lipidomics using traveling-wave ion mobility mass spectrometry. Nat Protoc 12, 797–813 (2017).

69. Wang, M. et al. Sharing and community curation of mass spectrometry data with Global Natural Products Social Molecular Networking. Nat Biotechnol 34, 828–837 (2016).

70. Ernst, M. et al. MolNetEnhancer: Enhanced Molecular Networks by Integrating Metabolome Mining and Annotation Tools. Metabolites 9, E144 (2019).

71. Morehouse, N.J. et al. Annotation of natural product compound families using molecular networking topology and structural similarity fingerprinting. Nature Communications 14, 308 (2023).

72. Pesic, A. et al. Champacyclin, a New Cyclic Octapeptide from Streptomyces Strain C42 Isolated from the Baltic Sea. Marine Drugs 11, 4834–4857 (2013).

73. Sekizawa, R. et al. Isolation and structural determination of phepropeptins A, B, C, and D, new proteasome inhibitors, produced by Streptomyces sp. J Antibiot (Tokyo) 54, 874–881 (2001).

74. Kanamori, Y., Iwasaki, A., Sumimoto, S. & Suenaga, K. Urumamide, a novel chymotrypsin inhibitor with a β-amino acid from a marine cyanobacterium Okeania sp. Tetrahedron Letters 57, 4213–4216 (2016).

75. Shishido, T.K. et al. Simultaneous Production of Anabaenopeptins and Namalides by the Cyanobacterium Nostoc sp. CENA543. ACS Chemical Biology 12, 2746–2755 (2017).

76. Aiken, S.G. et al. Iterative synthesis of 1,3-polyboronic esters with high stereocontrol and application to the synthesis of bahamaolide A. Nature Chemistry 15, 248–256 (2023).

77. Lysenkova, L.N., Turchin, K.F., Danilenko, V.N., Korolev, A.M. & Preobrazhenskaya, M.N. The first examples of chemical modification of oligomycin A. The Journal of Antibiotics 63, 17–22 (2010).

78. Chen, R. et al. Discovery of an Abundance of Biosynthetic Gene Clusters in Shark Bay Microbial Mats. Front Microbiol 11, 1950 (2020).

79. Scherlach, K. & Hertweck, C. Mining and unearthing hidden biosynthetic potential. Nat Commun 12, 3864 (2021).

80. Zhang, C. et al. Unexpected genetic and microbial diversity for arsenic cycling in deep sea cold seep sediments. NPJ Biofilms Microbiomes 9, 13 (2023).

81. Uritskiy, G.V., DiRuggiero, J. & Taylor, J. MetaWRAP—a flexible pipeline for genome-resolved metagenomic data analysis. Microbiome 6, 158 (2018).

82. Parks, D.H., Imelfort, M., Skennerton, C.T., Hugenholtz, P. & Tyson, G.W. CheckM: assessing the quality of microbial genomes recovered from isolates, single cells, and metagenomes. Genome Res 25, 1043–1055 (2015).

83. Olm, M.R., Brown, C.T., Brooks, B. & Banfield, J.F. dRep: a tool for fast and accurate genomic comparisons that enables improved genome recovery from metagenomes through de-replication. The ISME Journal 11, 2864–2868 (2017).

84. Chaumeil, P.A., Mussig, A.J., Hugenholtz, P. & Parks, D.H. GTDB-Tk v2: memory friendly classification with the genome taxonomy database. Bioinformatics 38, 5315–5316 (2022).

85. Letunic, I. & Bork, P. Interactive Tree Of Life (iTOL) v5: an online tool for phylogenetic tree display and annotation. Nucleic Acids Res 49, W293–W296 (2021).

86. Katoh, K. & Standley, D.M. A simple method to control over-alignment in the MAFFT multiple sequence alinment program. Bioinformatics 32, 1933–1942 (2016).

87. Capella-Gutierrez, S., Silla-Martinez, J.M. & Gabaldon, T. trimAl: a tool for automated alignment trimming in large-scale phylogenetic analyses. Bioinformatics 25, 1972–1973 (2009).

88. Nguyen, L.T., Schmidt, H.A., von Haeseler, A. & Minh, B.Q. IQ-TREE: a fast and effective stochastic algorithm for estimating maximum-likelihood phylogenies. Mol Biol Evol 32, 268–274 (2015).

89. Miller, M.A., Pfeiffer, W. & Schwartz, T. in 2010 Gateway Computing Environments Workshop (GCE) 1–8 (2010).

90. Jumper, J. et al. Highly accurate protein structure prediction with AlphaFold. Nature 596, 583–589 (2021).

91. Sullivan, M.J., Petty, N.K. & Beatson, S.A. Easyfig: a genome comparison visualizer. Bioinformatics 27, 1009–1010 (2011).

92. Kopylova, E., Noe, L. & Touzet, H. SortMeRNA: fast and accurate filtering of ribosomal RNAs in metatranscriptomic data. Bioinformatics 28, 3211–3217 (2012).

93. Patro, R., Duggal, G., Love, M.I., Irizarry, R.A. & Kingsford, C. Salmon provides fast and bias-aware quantification of transcript expression. Nat Methods 14, 417–419 (2017).

94. Zhao, Y., et al. TPM, FPKM, or Normalized Counts? A Comparative Study of Quantification Measures for the Analysis of RNA-seq Data from the NCI Patient-Derived Models Repository. Journal of Translational Medicine 19, 269 (2021).

95. Mohimani, H. et al. Dereplication of peptidic natural products through database search of mass spectra. Nat Chem Biol 13, 30–37 (2017).

96. Wandy, J. et al. Ms2lda.org: web-based topic modelling for substructure discovery in mass spectrometry. Bioinformatics 34, 317–318 (2018).

97. van der Hooft, J.J., Wandy, J., Barrett, M.P., Burgess, K.E. & Rogers, S. Topic modeling for untargeted substructure exploration in metabolomics. Proc Natl Acad Sci U S A 113, 13738–13743 (2016).

98. da Silva, R.R. et al. Propagating annotations of molecular networks using in silico fragmentation. PLoS Comput Biol 14, e1006089 (2018).

99. Djoumbou Feunang, Y., et al. ClassyFire: automated chemical classification with a comprehensive, computable taxonomy. J Cheminform 8, 61 (2016).

100. Shannon, P. et al. Cytoscape: a software environment for integrated models of biomolecular interaction networks. Genome Res 13, 2498–2504 (2003).

101. Dixon, P. VEGAN, a package of R functions for community ecology. Journal of Vegetation Science 14, 927–930 (2003).

