## Supplementary Figures for "A vast repertoire of secondary metabolites influences community dynamics and biogeochemical processes in cold seeps"

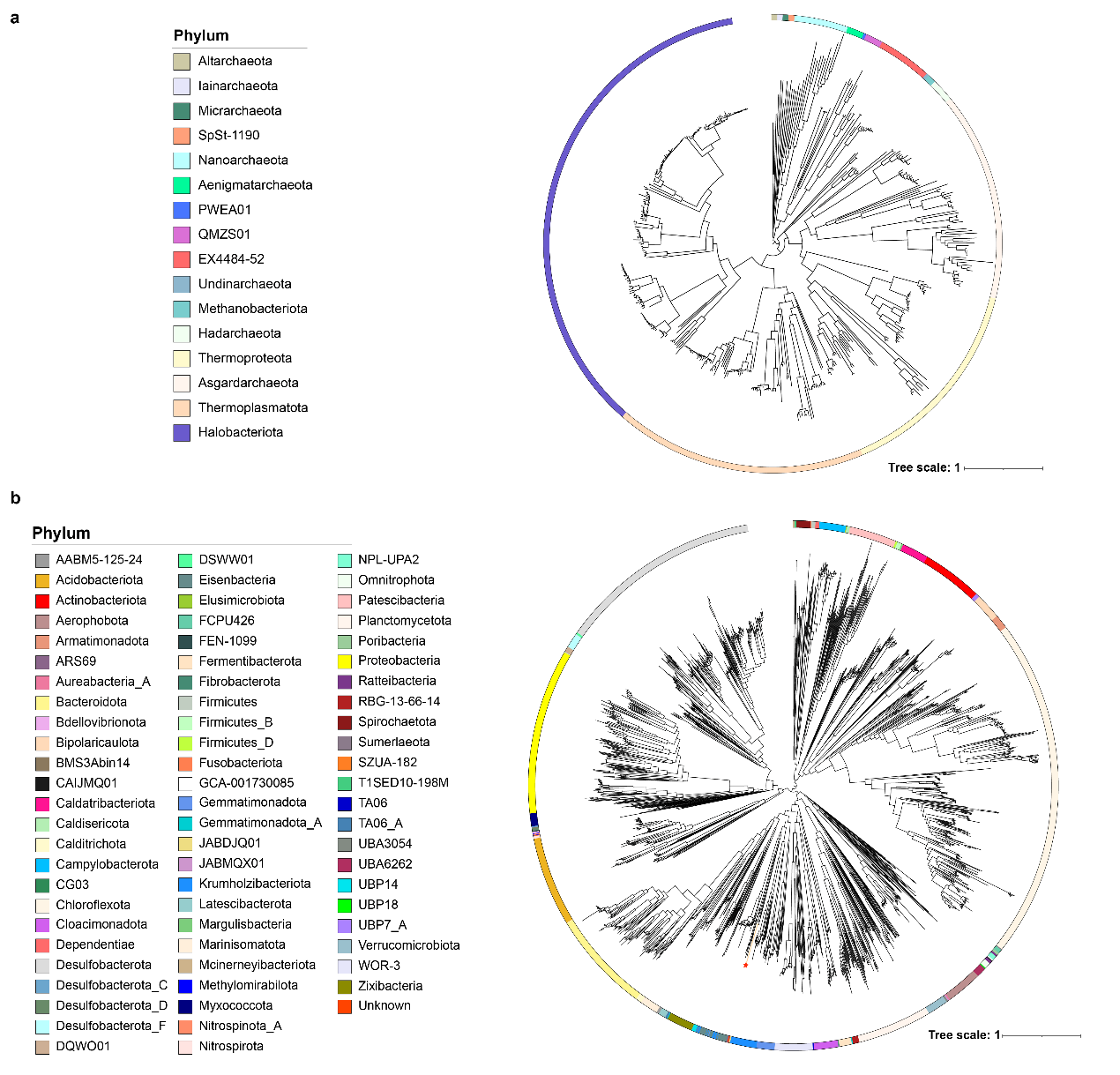

**Supplementary Figure 1. Maximum-likelihood phylogenetic trees of (a) archaeal and (b) bacterial MAGs recovered from deep sea cold seep sediments.**

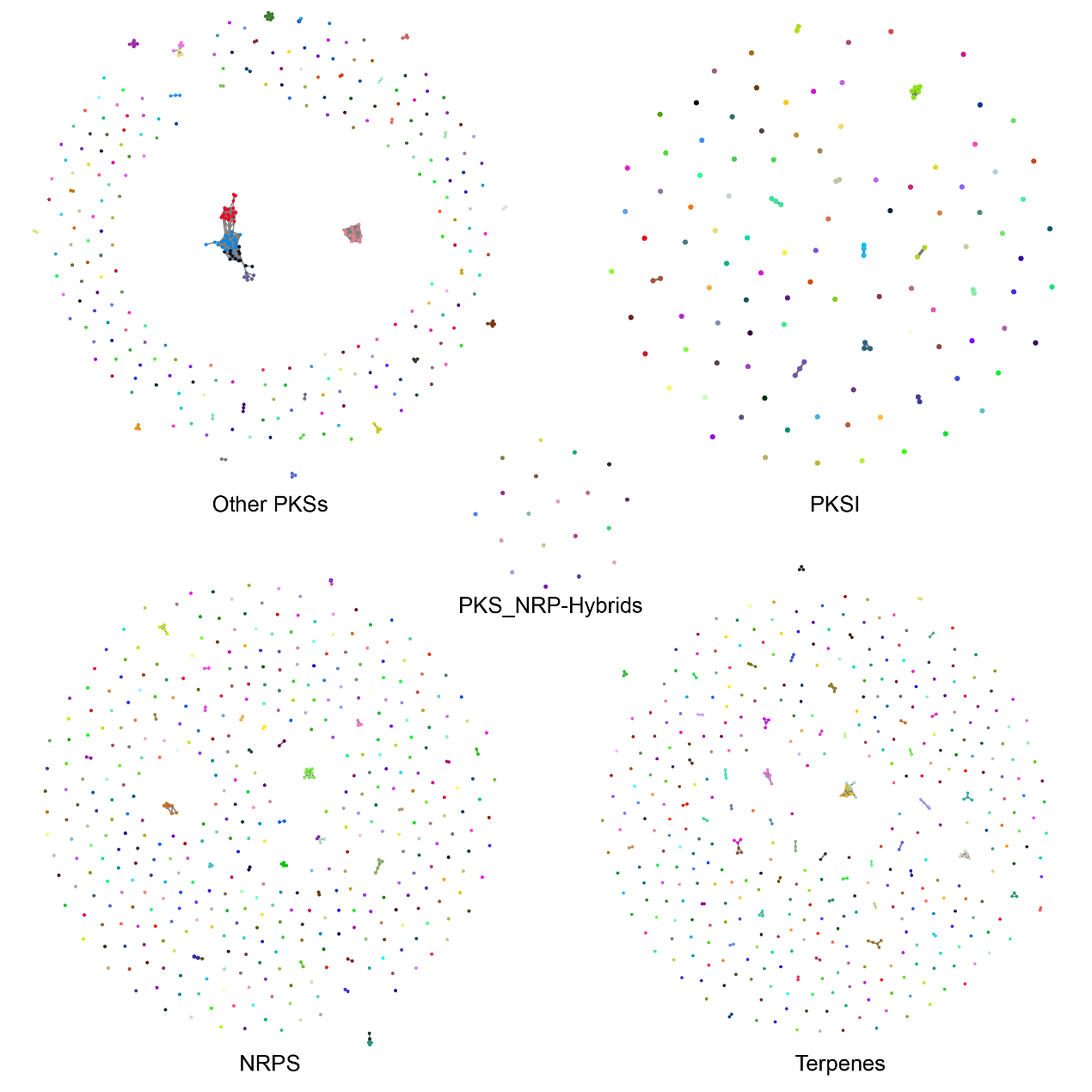

**Supplementary Figure 2. Network of the other five biosynthetic gene cluster types.** Their edges connect clusters that share genes, which are divided into five parts, including “Other PKSs”, “PKSI”, “PKS-NRP Hybrids”, “NRPS”, and “Terpenes”. The similarity networks are generated by BiG-SCAPE.

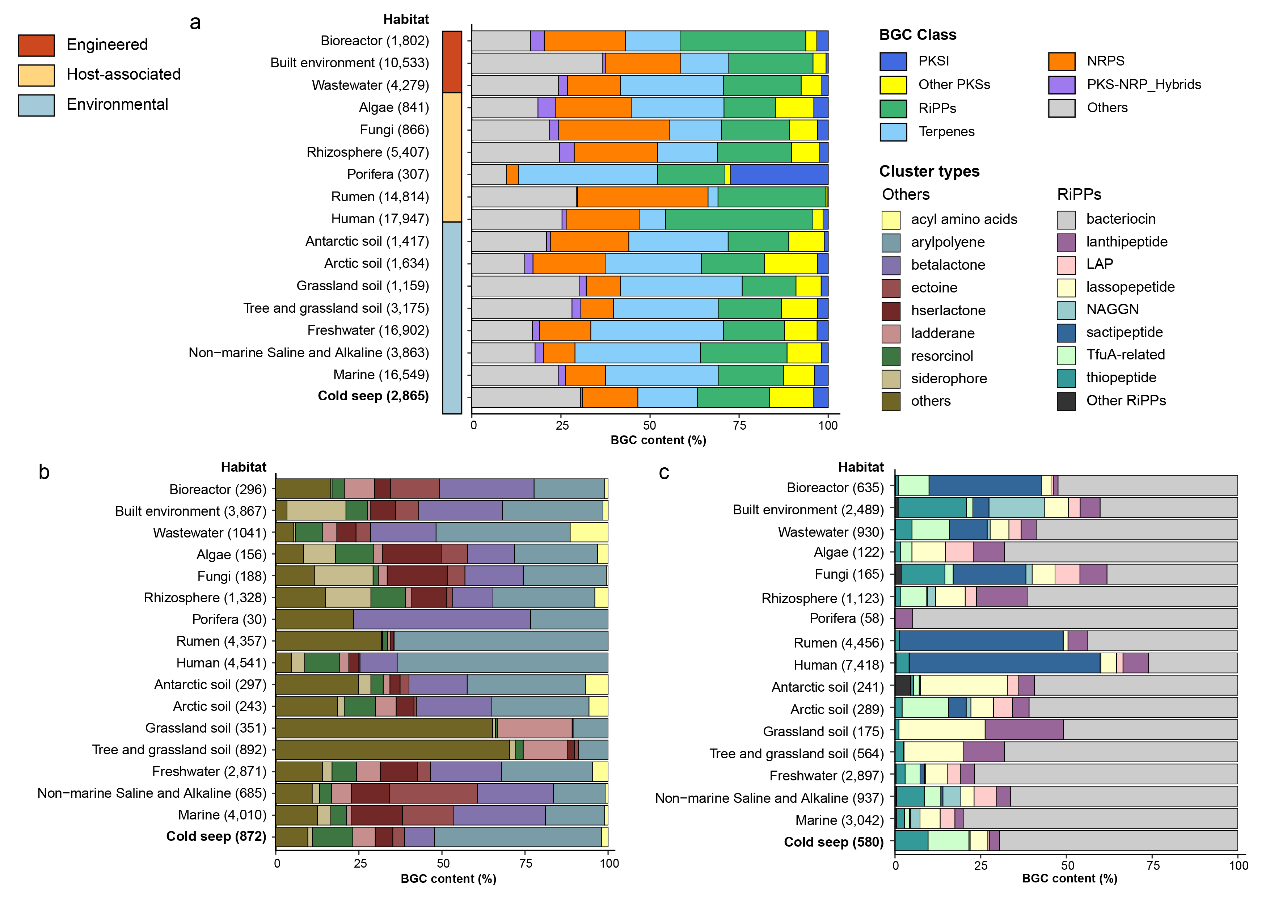

**Supplementary Figure 3. Relative frequencies of BGC types across different habitats.** **(a)** Relative proportions of seven BGC types across habitats. **(b)** Relative proportions of the Others class across habitats. **(c)** Relative proportions of the RiPPs class across habitats.

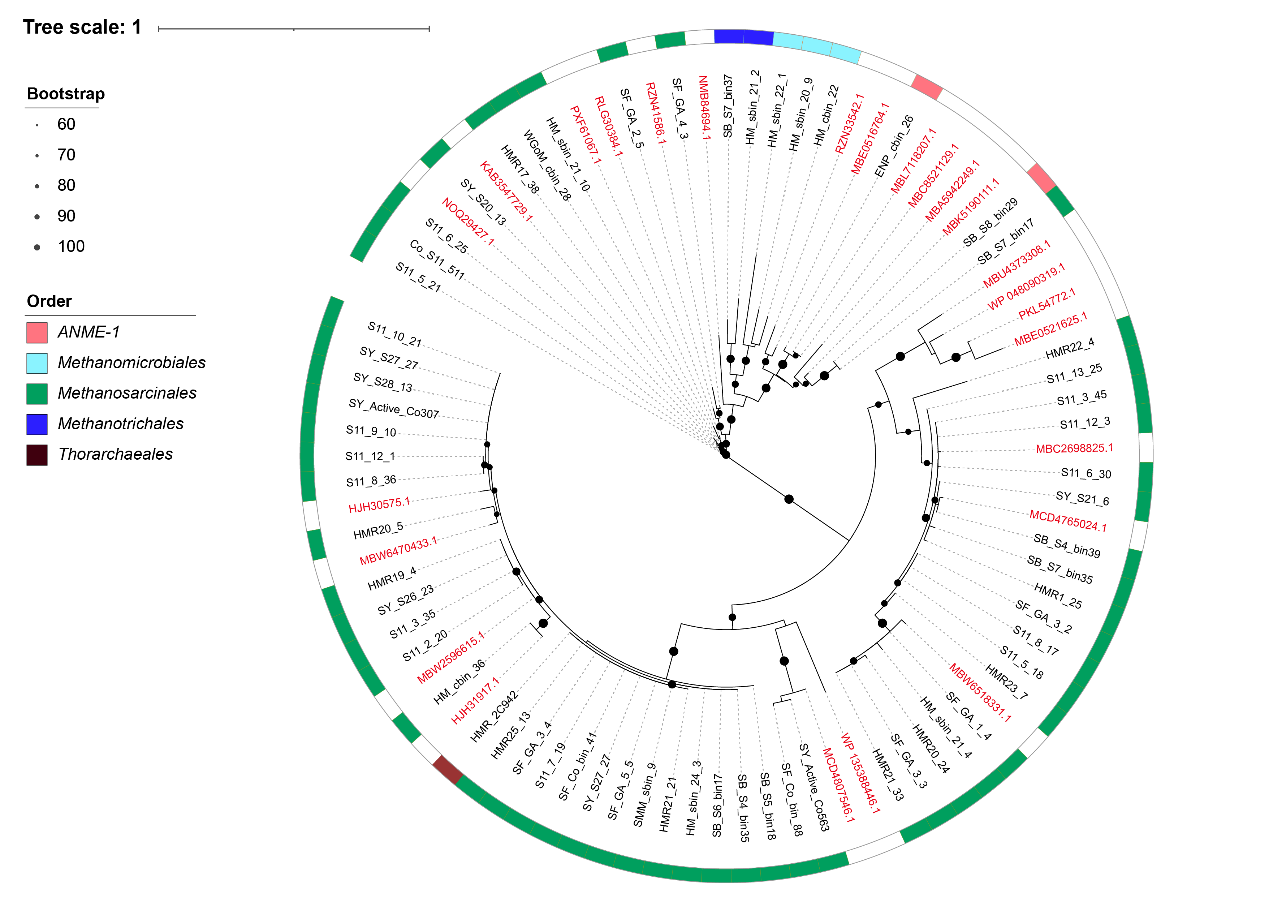

**Supplementary Figure 4. Maximum-likelihood phylogenetic tree of cold seep TfuA-like proteins.** Marked in red: 25 reference sequences (TfuA-related McrA-glycine thioamidation protein).

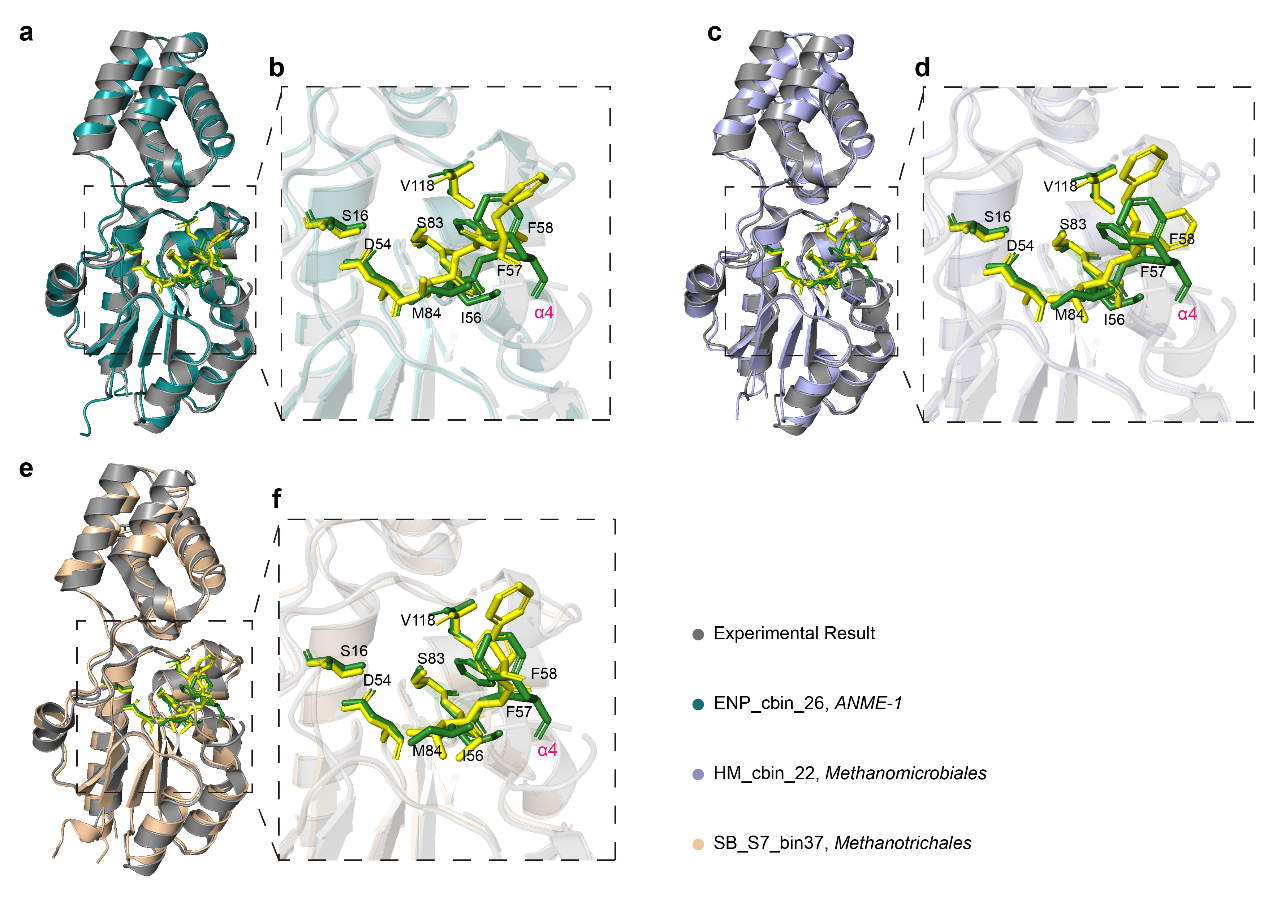

**Supplementary Figure 5.** **Structural comparison between three predicted TfuA-related McrA-glycine thioamidation proteins and the experimentally determined structure (6XPB, the crystal structure of TfuA involved in peptide backbone thioamidation from *Methanosarcina acetivorans*). (a, c, e)** The overall structural comparisions, showing the unique di-domain fold. The presumptive active site residues of experimental and predicted structures are shown as green and yellow sticks, respectively. **(b, d, f)** Close-up view of the putative active sites. Residues predicted to form the ThiS-binding pocket, and the α-helix that is implicated to mediate interactions with YcaO is marked as α4.

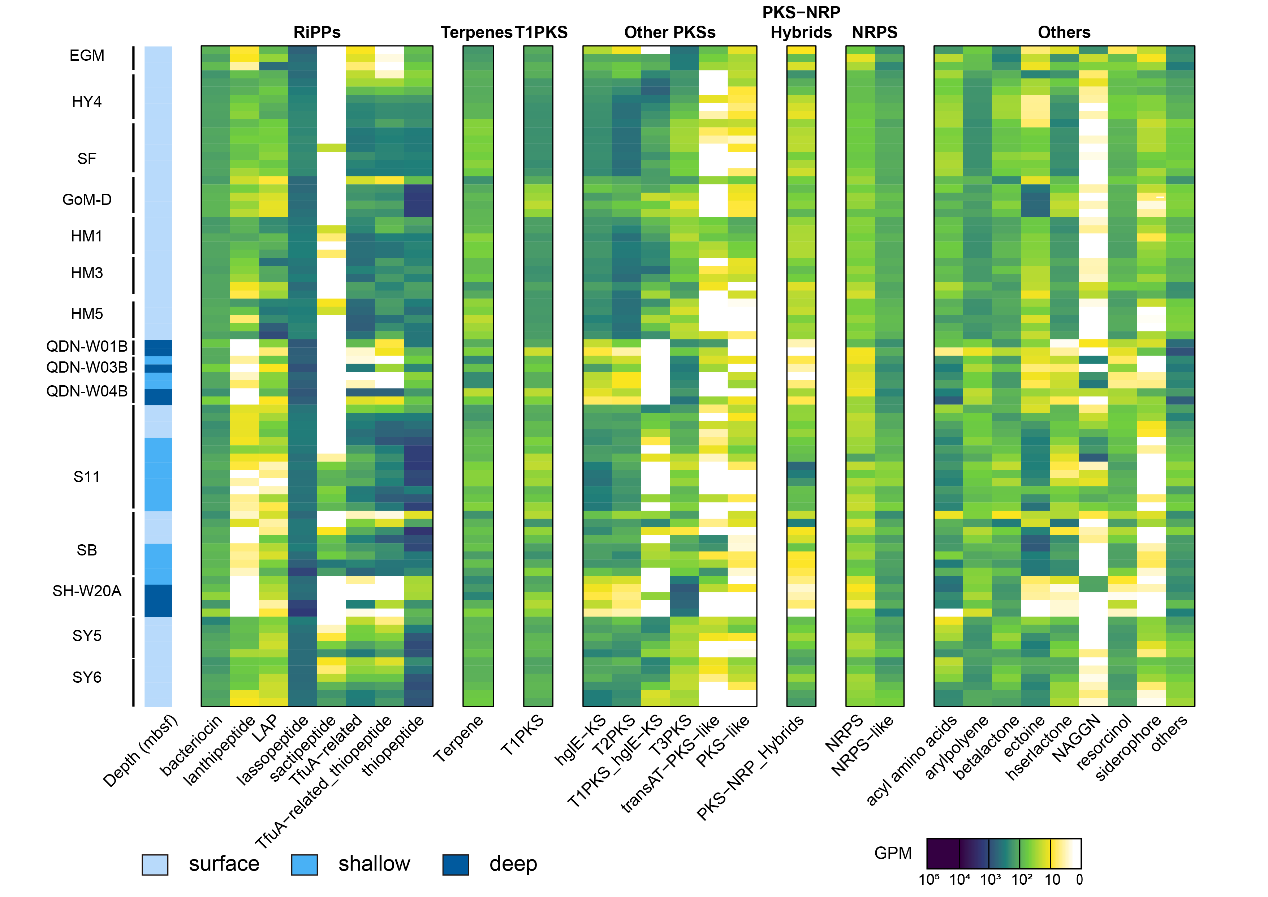

**Supplementary Figure 6.** **The heatmap shows the average abundance of BGCs among different BGC types at different sites.** Sediment depths are grouped as: surface, <1 mbsf; shallow, 1–10 mbsf; deep, >10 mbsf. BGC abundances are represented in the units of genes per million (GPM). Source data is available in **Supplementary Table 4**.

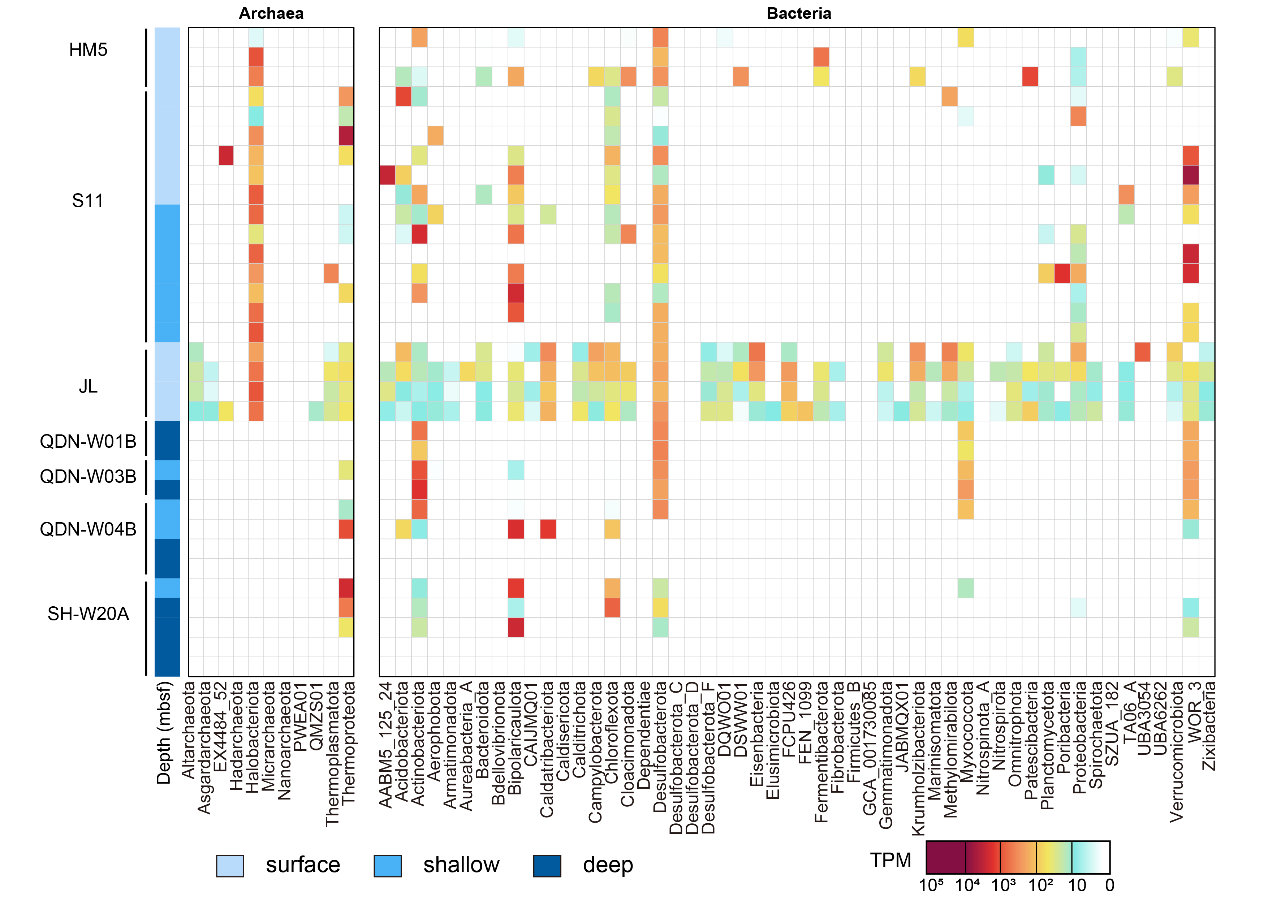

**Supplementary Figure 7.** **The heatmap shows the average transcript abundance of BGCs for each phylum.** 33 metatranscriptomic samples include the Haima cold seep, Jiaolong cold seep, Qiongdongnan basin, and Shenhu area. Transcript abundances are represented in the units of transcripts per million (TPM). Sediment depth are grouped into: surface, <1 mbsf; shallow, 1–10 mbsf; deep, >10 mbsf**.** Source data is available in **Supplementary Table 5**.

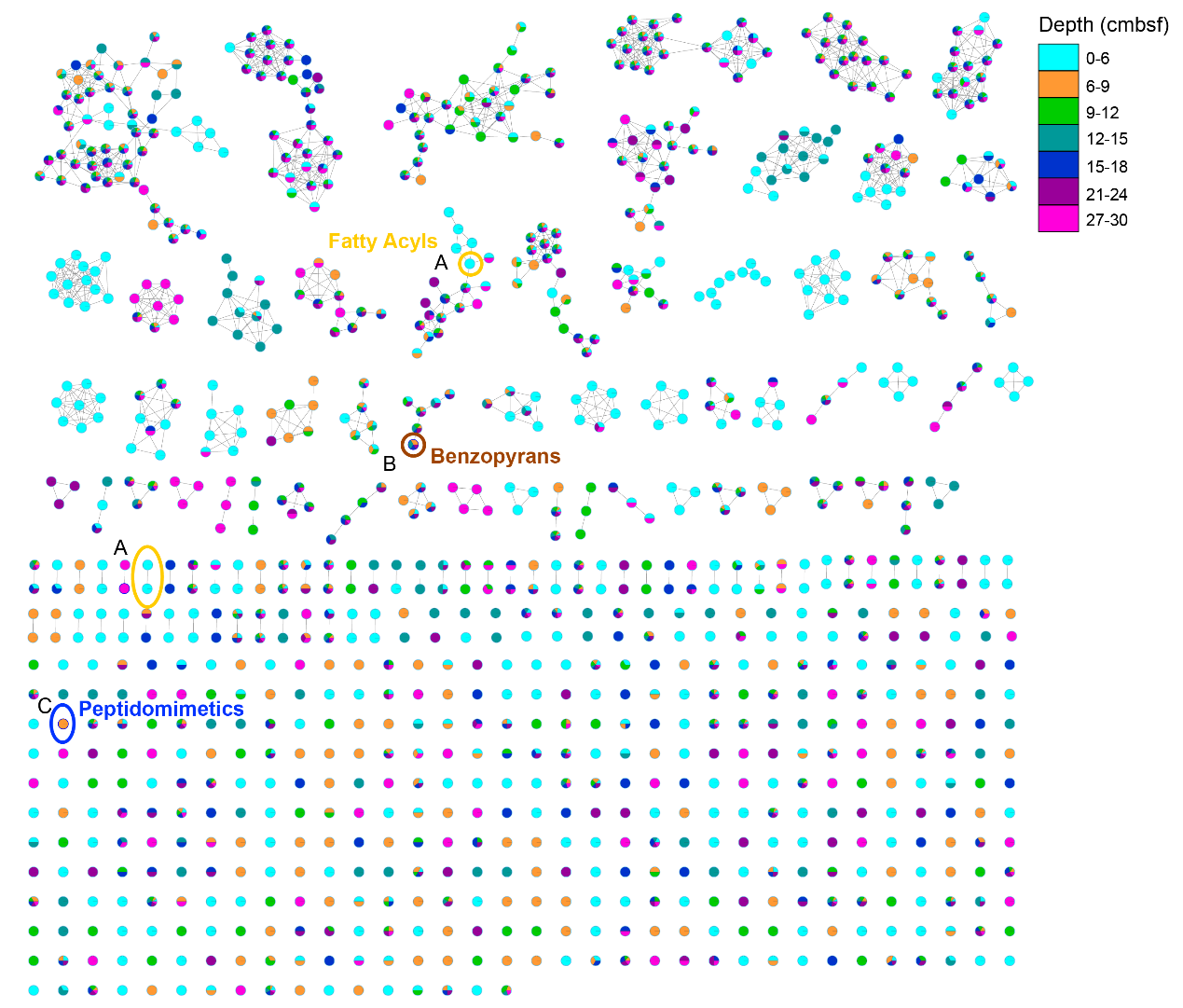

**Supplementary Figure 8.** **Molecular network of the hydrophilic metabolites in Qiongdongnan cold seep sediments.** Each dot is linked together to form clusters according to MS/MS spectrometry, colored by different depths. Five annotated features belong to three classes: benzopyrans, fatty acyls, and peptidomimetics by MolNetEnhancer. Detailed hydrophilic features are provided in **Supplementary Tables 6 and 8**.

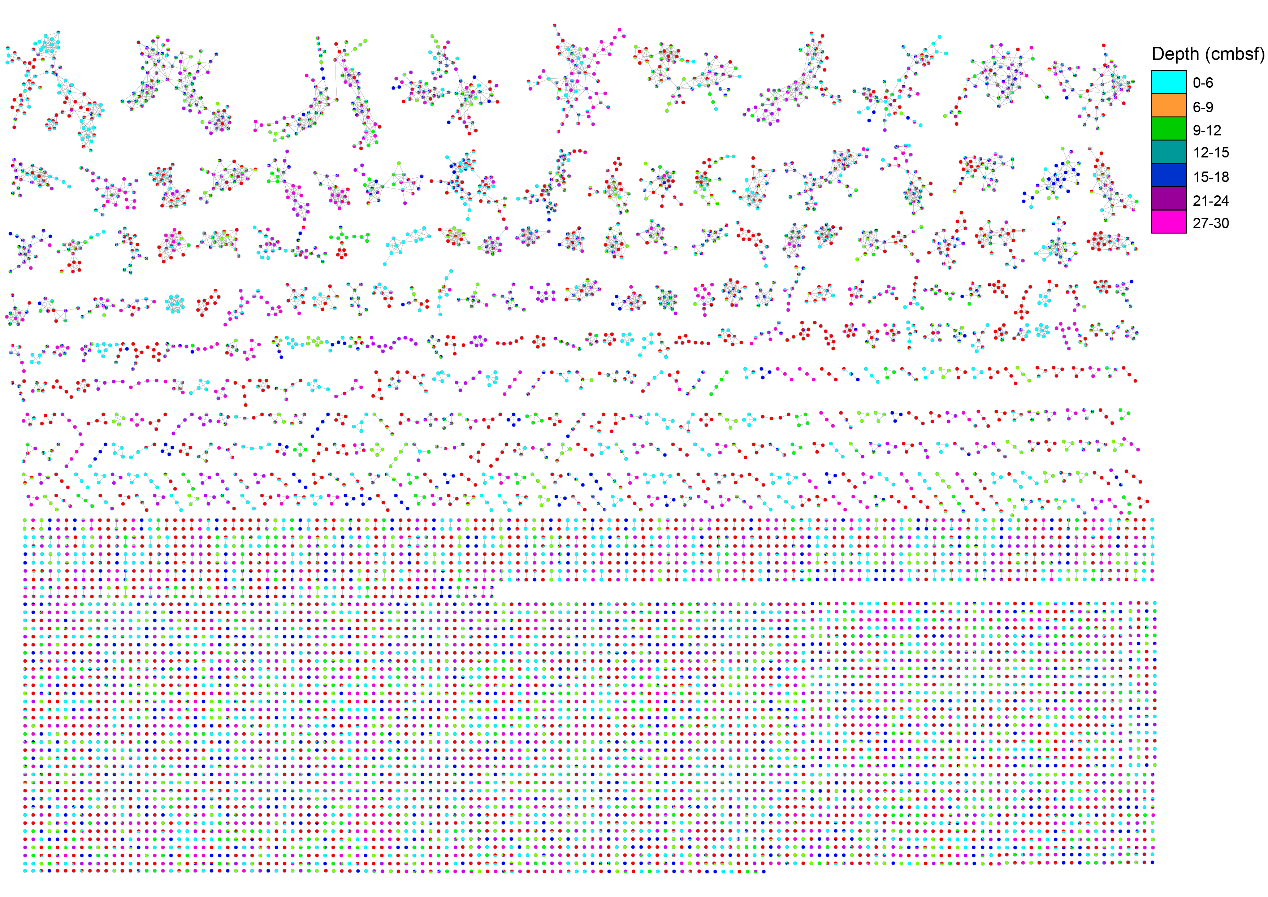

**Supplementary Figure 9.** **Molecular** **network of the lipophilic metabolites in Qiongdongnan cold seep sediments.** Each dot is linked together to form clusters according to MS/MS spectrometry, colored by different depths. Detailed nonpolar features are provided in **Supplementary Tables 7 and 9.**

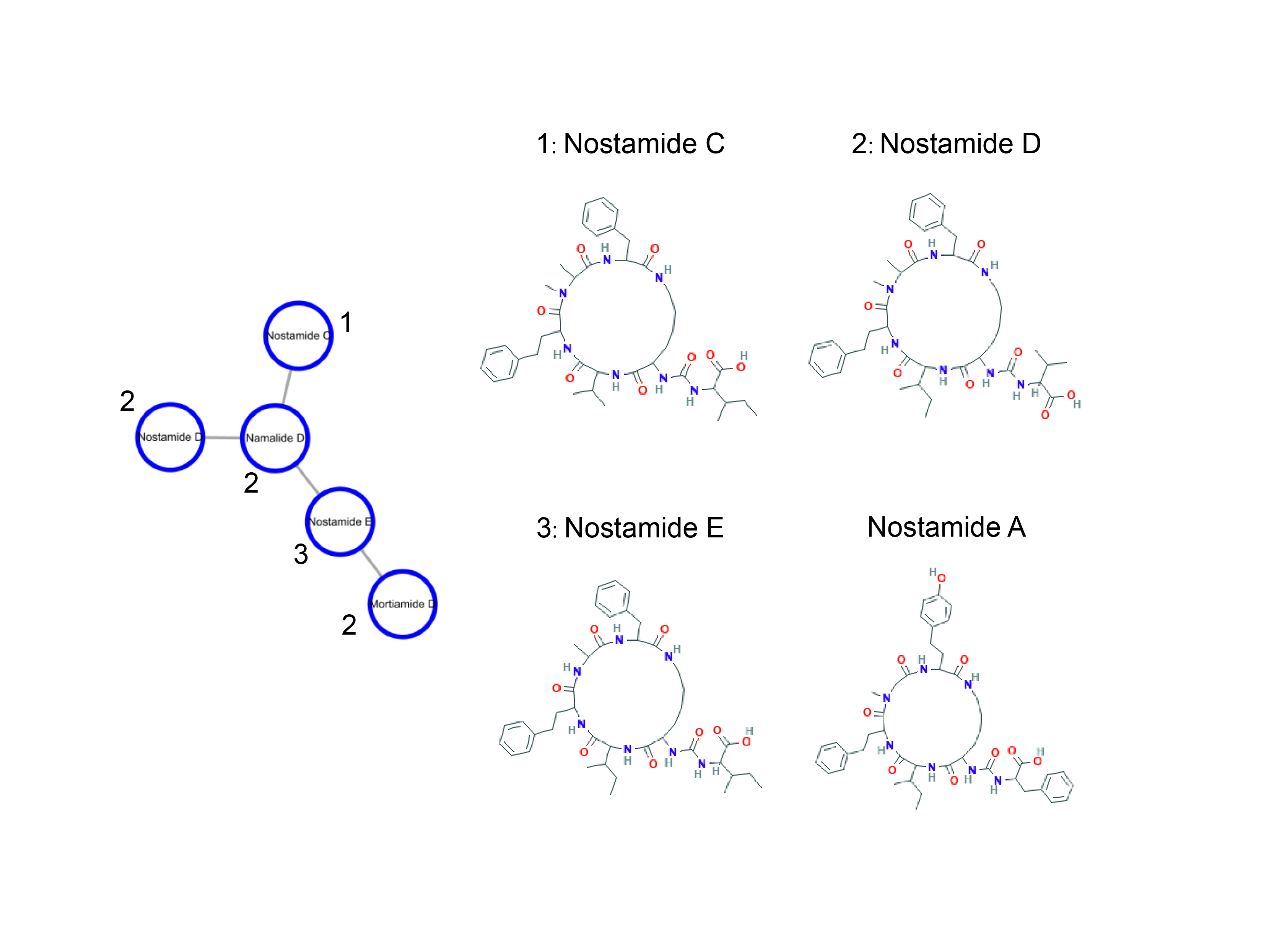

**Supplementary Figure 10.** **The cluster were annotated as nostamides by SNAP-MS (nostamide B, nostamide C, and nostamide E).** Nostamide A was predicted procduct based on 16 NRPS BGCs. Detailed data are in **Supplementary Table 9**.
